# LHPP expression in triple-negative breast cancer promotes tumor growth and metastasis by modulating the tumor microenvironment

**DOI:** 10.1101/2024.04.19.590151

**Authors:** Jeffrey Reina, Queralt Vallmajo-Martin, Jia Ning, Aubrey N. Michi, Kay Yeung, Geoffrey M. Wahl, Tony Hunter

## Abstract

Triple-negative breast cancer (TNBC) is a highly aggressive and metastatic form of breast cancer that lacks an effective targeted therapy. To identify new therapeutic targets, we investigated the phosphohistidine phosphatase, LHPP, which has been implicated in the development of several types of cancer. However, the full significance of LHPP in cancer progression remains unclear due to our limited understanding of its molecular mechanism. We found that levels of the LHPP phosphohistidine phosphatase were significantly increased in human breast cancer patients compared to normal adjacent tissues, with the highest levels in the TNBC subtype. When LHPP was knocked out in the MDA-MB-231 human TNBC cell line, cell proliferation, wound healing capacity, and invasion were significantly reduced. However, LHPP knockout in TNBC cells did not affect the phosphohistidine protein levels. Interestingly, LHPP knockout in MDA-MB-231 cells delayed tumor growth and reduced metastasis when orthotopically transplanted into mouse mammary glands. To investigate LHPP’s role in breast cancer progression, we used next-generation sequencing and proximity-labeling proteomics, and found that LHPP regulates gene expression in chemokine-mediated signaling and actin cytoskeleton organization. Depletion of LHPP reduced the presence of tumor-infiltrating macrophages in mouse xenografts. Our results uncover a new tumor promoter role for LHPP phosphohistidine phosphatase in TNBC and suggest that targeting LHPP phosphatase could be a potential therapeutic strategy for TNBC.

## Introduction

Protein phosphorylation is a crucial molecular process that regulates many cellular functions, including cell growth, division, differentiation, and apoptosis (1). During phosphorylation a phosphate group is added to a protein, typically on serine, threonine, or tyrosine residues, generating the three “classical” phospho-amino acids, which have been extensively studied and long implicated in cancer progression (1). However, 6 other amino acids can be phosphorylated, including histidine and these have been historically overlooked (2). The 6 non-canonical phospho-amino acids are heat and acid labile, rendering routine molecular assays more challenging, thus we understand little about biological consequences of this type of phosphorylation (2). In 2015, to address a lack of tools to study histidine phosphorylation, our group developed monoclonal antibodies against the two isoforms of phosphohistidine (pHis). Using these antibodies, we can detect pHis levels on either of its 1- and 3-phosphoisomers by conventional methods such as immunoblot, immunofluorescence, and mass spectrometry (3). Recently, the histidine phosphorylation field has gained momentum due to the discoveries of its involvement in the development and progression of liver cancer (4) and neuroblastoma (5). In the context of phosphorylation, numerous studies highlight the dysregulation of protein kinases and phosphatases in various types of cancer, which are the enzymes that add or remove a phosphate group on proteins (1). Therefore, the study of histidine kinases and phosphatases is crucial to understanding how histidine phosphorylation is regulated (6, 7). Currently, there are only two well-studied histidine kinases: NME1 and NME2, and three known phosphohistidine phosphatases: PHPT1, PGAM5, and LHPP (2, 7). Whether phosphohistidine or any histidine kinase or phosphohistidine phosphatase is involved in other cancer types remains largely unknown, including in breast cancer.

Breast cancer is prevalent among women worldwide, with triple-negative breast cancer (TNBC) being a highly aggressive and metastatic subtype lacking efficient targeted therapy options (8, 9). Therefore, a deeper understanding of histidine kinases and phosphatases is essential to identify potential therapeutic targets for breast cancer treatment. For instance, NME1, a histidine kinase, was initially linked to the low metastatic potential of melanoma cells and was subsequently identified as a metastasis suppressor (10). Additionally, its high expression in primary tumors of the liver and colon was observed to decrease during metastasis (11). PHPT1 has been reported to be upregulated in lung cancer patients and cell lines (12, 13), associated with decreased proliferation in hepatocellular carcinoma (14), and increased levels have been linked to lower overall survival in clear-cell RCC patients (15). PGAM5, on the other hand, is upregulated in liver cancer and its knockout inhibits the proliferation, migration, and invasion of gastric cancer cells (16, 17). Only the histidine kinase NME1 has been studied in breast cancer, where it was shown to inhibit the migration of breast cancer cell lines (18). In this study, we focused on the role of another pHis-associated enzyme, the phosphohistidine phosphatase LHPP which was first described to be critical in liver cancer (4), yet both its molecular mechanism and role in breast cancer require further investigation.

LHPP, also known as phospholysine phosp<u>h</u>ohistidine inorganic <u>p</u>yrophosphate <u>p</u>hosphatase, remains somewhat of a biological enigma as its cellular functions are largely unknown. It has been shown to inhibit tumor progression in various types of cancers including cervical, pancreatic, colorectal, gastric, renal, glioblastoma, bladder and thyroid cancers (19–27). A role for LHPP was first explored in liver cancer, where it was shown to inhibit autochthonous hepatocellular tumor growth driven by combined loss of Pten and Tsc1, suggesting that LHPP acts as a tumor suppressor in this context (4). Since that report, several studies on other cancer types have followed (27, 28). In liver cancer, renal cell carcinoma, prostate cancer, glioblastoma, colorectal cancer, pancreatic cancer, cervical cancer, melanoma, bladder cancer, thyroid cancer, and non-small cell lung cancer, LHPP expression was associated with the inhibition of cell proliferation, induction of apoptosis, and suppression of metastasis in cancer cell lines and mouse models (27).

However, the molecular mechanism by which LHPP regulates tumor cell growth is still poorly understood, and thus far, no LHPP phosphohistidine protein substrates have been identified. In this study, we investigated the role of LHPP phosphohistidine phosphatase in triple-negative breast cancer (TNBC), its impact on tumor growth and metastasis, and evaluated the changes in cell signaling pathways by altering LHPP expression levels. We discovered that LHPP protein levels are significantly increased in TNBC patient tumors, compared to normal breast tissue, and that genetic loss of LHPP delays tumor growth and reduces metastasis of TNBC cell lines in mouse xenografts by modulating the tumor microenvironment.

## Methods

### Human samples

Human tissue microarrays of formalin-fixed paraffin-embedded (FFPE) breast tumors and normal patient tissues were obtained from Tissuearray.com (Biomax, Inc.). The slides analyzed in this study were #BR1503f, #BR1202a, and #BR1401. To classify the different tumors by subtype, the expression of estrogen receptor (ER), progesterone receptor (PR), and HER2 receptor was used, as provided by Tissuearray.com (Biomax, Inc.). The Biorepository and Tissue Technology Shared Resources at Moores Cancer Center, University of California San Diego (UCSD), provided all de-identified human primary breast tumor and normal tissue samples, which had been previously collected (see Supplementary methods).

### Immunohistochemistry (IHC)

Mouse tissue was fixed in 4% paraformaldehyde overnight and submitted to the Tissue Technology Shared Resource – Histology Core at UCSD for paraffin embedding and sectioned to 5 µm. Mouse and human FFPE tissues were deparaffinized in xylene and rehydrated through graded ethanol solutions (100%, 95%, 70% and 50% EtOH). Antigen retrieval was performed at 95-100°C in 10 mM sodium citrate buffer with 0.05% Tween-20, pH 6.0, for 15 min. Endogenous peroxidases were quenched using 3% H_2_O_2_ for 10 min and blocked in 5% BSA, 5% normal goat serum, 0.1% Tween-20 in TBS for 1 h. Primary antibodies were diluted in blocking buffer and incubated overnight at 4°C in a humidified chamber. After washing with 0.1% TBS-T, the slides were incubated with SignalStain Boost IHC Detection Reagent (Cell Signaling Technology, 8114) for 30 min at room temperature. After washing three times with 0.1% TBS-T, the signal was developed using ImmPACT DAB Substrate Kit (Vector Laboratories, SK4105). Slides were counterstained with hematoxylin and then dehydrated through reverse-graded ethanol solutions (50%, 70%, 95%, and 100% EtOH), cleared in xylene, and mounted with DPX mounting media (Millipore Sigma, 317616). The following primary antibodies were used with the indicated dilutions: rabbit anti-LHPP (Sigma Millipore ZRB1050, 1:250), rabbit anti-human vimentin (Abcam ab8069, 1:1000), rabbit anti-F4/80 (Cell Signaling Technology, 70076, 1:200), and rabbit anti-Ki67 (Abcam, ab15580, 1:50).

### Images acquisition and quantification of IHC staining

Immunohistochemistry images were acquired using a VS120-L100 Virtual Slide System (Olympus) at 40X objective using the same scanning parameters between slides. For quantification of the DAB staining, QuPath Software (v.0.4.4) was used. The stains were automatically deconvoluted by using the estimates vector stains function for each antibody. Subsequently, the positive cell function was used to count the number of cells that were positively and negatively stained. Cells were automatically detected by the software using nuclear hematoxylin staining. Three intensity thresholds were set for DAB-positive cells, threshold 1+ was set at 0.2 OD, threshold 2+ at 0.3 OD, and threshold 3+ at 0.4 OD. For the score compartment, the cell DAB OD maximum parameter was used. The rest of the settings were kept as default, and the same settings were used for all images stained with the same antibody. To measure both the intensity and percentage of cells stained, an H-score was used and calculated automatically by QuPath software. Briefly, the H-score is determined by the sum of multiplying the percentage of cells stained 1+ by 1, the percentage of cells stained 2+ by 2, and the percentage of cells stained 3+ by 3 giving a score from 0 to 300 (29).

### Kaplan-Meier survival plots

The Kaplan-Meier Plotter online tool (www.kmplot.com) was used to obtain overall survival curves for breast cancer patients expressing high and low LHPP. Kmplotter uses curated mRNA-sequencing data from GEO, EGA, and TGCA (30). To obtain the survival curve of ER^+^PR^+/-^ HER2^+/-^ patients, in Kmplotter the categories ER status was indicated positive. To obtain the survival curve of ER^-^PR^-^HER2^+^ patients, in Kmplotter the categories ER, and PR status were indicated negative and HER2 status positive. To obtain the survival curve of TNBC patients, in Kmplotter the categories ER, PR, and HER2 status were indicated negative.

### Immunoblotting

Cell lysates were prepared using 1X RIPA buffer (50 mM Tris-HCl pH 8.8, 1% TritonX-100, 0.1% SDS, 0.5% sodium deoxycholate, 150mM NaCl, 1mM EDTA) supplemented with Mini Complete EDTA-free protease inhibitor (Roche, 11836170001) and PhosSTOP (Roche, 4906837001). Samples were probe sonicated using three bursts of 20% amplitude until 40J was reached and incubated for 1 h at 4°C. Samples were centrifuged at 15,000 x g for 15 min to collect soluble protein. Protein concentration was quantified using a BioRad DC Protein Assay (Bio-Rad, 500-0116). Whole-cell lysates were separated by either 12.5% or 4-12% SDS-PAGE Bis-Tris gels (Invitrogen) and transferred to a 0.45 um PVDF membrane overnight at 4°C. Membrane was blocked in 0.2X PBS with 0.1% casein (Bio-Rad, 1610783) for 1 h and primary antibodies were incubated for 2 h at RT or 4°C overnight in blocking buffer with 0.1% Tween-20. Membranes were washed three times for 5 minutes in 0.1% TBS-T and incubated with goat anti-rabbit IgG secondary antibody, Dylight800 (Invitrogen SA535571, 1:20,000), and goat anti-mouse AlexaFluor680 (Invitrogen A-21058, 1:20,000) diluted in blocking buffer with 0.1% Tween-20 and 1% SDS. The following antibodies were used at the indicated dilution: rabbit anti-LHPP (Sigma Millipore, ZRB1050, 1:1000), Mouse β -actin (Millipore Sigma, A1978, 1:1000), streptavidin AlexaFluor680 (Invitrogen, S32358, 1:20,000), mouse anti-Flag tag (Millipore Sigma, F3165, 1:5000). Membranes were scanned on a LI-COR Odyssey Imager, and image quantification was performed in Image Studio version 2.1.10 (LI-COR). For phosphohistidine immunoblotting, all buffers were adjusted to pH 8.8, and a heat control of each sample was boiled at 95°C for 15 min, in which genuine phosphohistidine signals should diminish, given its heat instability. The gel separation and membrane blocking were performed at 4°C. The primary antibodies were incubated 1 h at 4°C and 1 h at room temperature. The rest of the procedure was performed at room temperature. For primary antibodies, the 3-phosphohistidine SC44-1 antibody (0.5 µg/ml) and the 1-phosphohistidine SC1-1 antibody (0.5 µg/ml) isolated from hybridoma cell culture medium in our lab were used.

### Cell lines

MDA-MB-231 human breast cancer cell line was obtained from the Salk Cancer Center collection. Cells were cultured in DMEM (Gibco, 10-013-CV) and supplemented with 10% fetal bovine serum (Omega Scientific, FB-01) and 1% penicillin/streptomycin (Gibco, 15140122) at 37°C with 5% CO_2_. Cells were fed every 48 h and were passaged at 80% confluence.

### Generation of CRISPR Cas9 knockout LHPP cell line

The plasmid px330 expressing mCherry was used as a backbone to clone the sgRNA CACATCCAACCCAAACTGTGO that targets human LHPP exon 3. This sgRNA was designed using Benchling Biology Software (https://benchling.com), where the top 4 sgRNAs with the highest on-target score were tested. Once the sgRNA was inserted into the px330 plasmid, it was transiently transfected into MDA-MB-231 cells using Lipofectamine 3000. After 48 h the mCherry-positive cells were sorted at Salk’s Flow Cytometry Core and Hoechst 33342 (Invitrogen, H3570) staining was used to determine cell viability. The cells were single-sorted into a 96-well plate and the remainder were grown in limited dilution conditions on 10 cm plates to obtain single clones. The clones were expanded and screened for LHPP expression by both Sanger sequencing and immunoblotting.

### Cell proliferation assay

MDA-MB-231 cell proliferation was assessed by seeding 10,000 cells per well into 96 well plates and imaged every 4 h using the IncuCyte ZOOM^®^ (Essen Bioscience) for 72 h. Confluence and cell growth readings were performed using Incucyte™ Software using phase-contrast images.

### Scratch wound assay

MDA-MB-231 cell scratch wound assay was performed by seeding 50,000 cells per well into an Incucyte^®^Imagelock 96-well plate (Essen Bioscience). At 100% confluence, a scratch wound was made using a WoundMaker™ (Essen Bioscience). Cells were washed twice in DMEM to remove cell debris and complete growth medium was added. The plate was imaged every 4 h for 72 h using the IncuCyte S3 v2019A system (Essen BioScience). Relative wound confluence was determined by using the Incucyte™ Scratch Wound Cell Migration Software Module.

### Spheroid formation

Spheroid formation of MDA-MB-231 cells was performed by seeding 10,000 cells per well on ultra-low attachment 96 well plates (Corning, 7007) and imaged on an inverted microscope after 72 h. The circularity and area of the spheroids were analyzed in ImageJ using a customized macro. Briefly, a Gaussian Blur filter with sigma=3 was used, then it was converted to a binary mask and the analyze particles function was used with a minimum size of 800 pixels. This script was run on all spheroid images from all cell types as a batch.

### Cell invasion

Cell culture plates for cell invasion assays were prepared by adding 150 μL of 1 mg/mL Cultrex Reduced Growth Factor Basement Membrane Extract (R&D Systems, 3433-005-R1) to a 24-well cell culture insert with an 8 μm pore size (Falcon, 353097). After incubating at 37°C for 30 min, 75,000 cells in 150 μL DMEM medium were added to each insert, and 800 μL of complete growth medium supplemented with 20% FBS was added to each well on a 24-well cell culture plate. After 24 h, cells were fixed using 100% methanol for 20 min at room temperature. Then the transwells were stained with 0.5% crystal violet in a 20% methanol solution for 20 min. The remaining crystal violet and non-invading cells on the transwells were removed with a cotton swab. Subsequently, the transwells were washed three times with distilled water for 5 min. Images were obtained and the cells were counted using ImageJ with a customized macro. Briefly, a binary mask was applied to the images, and the analyze particles function was used to count invading cells with a minimum size of 300 pixels and minimum circularity of 0.05 to not detect the scale bar. In experiments where the images were abundant with invading cells, a binary erode function was performed twice and the binary watershed function was utilized to separate the cells that were in close proximity to one another. A batch of the same script, with identical parameter settings, was executed on all invading cell images from the same experiment.

### Mice

All mouse experiments were performed with approval from the Animal Resources Department at the Salk Institute for Biological Studies and the Institutional Animal Care and Use Committee (IACUC), under the 2011-0005 approved protocol. CB17 SCID female mice were purchased from Charles River Laboratories. All mice were housed in a controlled environment, in a pathogen-free animal facility at the Salk Institute for Biological Studies, with a 12 h light-dark cycle at 23°C and 30-70% humidity, fed rodent chow *ad libitum,* and cared for by the Animal Research Department.

### Orthotopic mammary gland transplantation

CB17 SCID female mice between 10 and 13 weeks old were anesthetized, and a midline and angled lateral incision between the left nipple #4 and #5 was made. The skin flap was loosened from the abdominal body exposing the #4 mammary gland. In parallel, 1 x 10^6^ cells from MDA-MB-231 WT, LHPP KO clone 3 (KO3), and clone 11 (KO11) were prepared in a 7.5 μL suspension in PBS. Next, a cell suspension was mixed with 1 μL of trypan blue and 8.5 μL of growth factor reduced Matrigel (1:1, Corning 356231) in a total of 17 μL volume which was then injected into the #4 left mammary gland. Skin flaps were sutured, and mice were closely monitored during recovery. Tumor measurements were performed once per week until they reached a volume higher than 1000mm^3^, marking the study’s endpoint, in which mice were euthanized by CO_2_ asphyxiation and cervical dislocation, and tissues were collected.

### Tumor and tissue collection and dissociation

Mice were euthanized at the experimental endpoint and the tumors were dissected and weighed. Mammary glands (#2/3, #4 right, and #2/3 left), lungs, and livers were inspected for macrometastasis and dissected. All tissues were then rinsed in PBS, minced with a razor blade or scissors, and collected in 5 mL of dissociation media consisting of Epicult-B Basal medium containing Mouse Epicult B supplement (Stemcell Technologies, 05610), 5% FBS, 1% penicillin/streptomycin, 2.5 μg/mL Amphotericin B, Collagenase and Hyaluronidase (1650 U/550 U, Stemcell Technologies, 07912) and 4 μg of hydrocortisone (Sigma, 50237). Tissue samples were then incubated at 37°C with agitation for 3 h. To obtain single cells the tissue was centrifuged at 450 x *g* for 3 min and the cell pellet was resuspended in 1 mL Dispase 5 U (Stemcell Technologies, 07913) with 100 μL of DNase I 1 mg/mL (Sigma, 10104159001). The cells were centrifuged and resuspended in 1 mL 0.25% Trypsin. Subsequently, red blood cells were lysed using a 0.64% NH_4_Cl solution containing 10 mM EDTA pH 7.4-7.6. The cells were passed through a 100 µm cell strainer, centrifuged, and resuspended in 250-500 μL Hank’s balanced salt solution (Gibco, 14025092) containing 2% FBS, 10 μg DNase I, and 1 μg/mL Purified Rat Anti-Mouse CD16/CD32 (Mouse BD Fc Block^TM^, BD Biosciences 553142) and quantified by counting using trypan blue to exclude non-viable cells.

### Single-cell staining and Fluorescence-Activated Cell Sorting (FACS)

Cells were stained with lineage markers including CD45-BV421 (BioLegend, 103134), Ter119-BV421 (BioLegend, 116234), and CD31-BV421 (BioLegend, 102424), epithelial markers including CD326 (EpCAM)-Alexa Fluor 647 (BioLegend, 118212), CD49f-APC/Cy7 (BioLegend, 313628), and CD61-PE/Cy7 (BioLegend, 104318), and two human-specific markers HLA-ABC-FITC (BioLegend, 311404) and hCD44-PE (BioLegend, 338807), and counterstained with DAPI. Cells were then analyzed and sorted in a FACSAria^TM^ Fusion sorter, which was set to acquire and record 500,000 events for each sample. Additionally, single, live, lineage-negative, and human cells (HLA-ABC^+^/CD44^+^) from tumor samples were sorted, centrifuged, and lysed in RIPA 1X buffer. Flow cytometry analysis was performed using FlowJo (v.10.8.2) software.

### Messenger RNA sequencing (mRNA-seq)

Three 60 mm plates of MDA-MB-231 cells and LHPP KO (KO11 clone) MDA-MB-231 cells 70-90% confluent were washed two times with cold PBS and 600 μL TRIzol reagent (Invitrogen, 15596018) was used to collect the cells. Then, the Direct-zol RNA miniprep kit (Zymo Research, R2050) was used to extract total RNA. Samples were quantified and 1 mg of total RNA was submitted to Novogene Co. (Sacramento, CA, USA) for mRNA sequencing and bioinformatic analysis. Briefly, RNA sample quality control was performed using a Qubit, mRNA library preparation was performed using poly A enrichment, and NovaSeq PE150 (6 GB raw data per sample) was used to generate the reads. The reads were trimmed for low-quality or adapter-containing regions and mapped to the reference genome *Homo sapiens* (GRCh38/hg38). Gene expression quantification was performed and Fragments Per Kilobase of transcript sequence per Million base pairs sequenced (FPKM) (31) was calculated. Differential gene expression (DEG) was performed using DeSeq2 (32).

### Generation of LHPP-TurboID and TurboID stable cell lines

The pLVX, pLVX-TurboID-Flag tag, and pLVX-LHPP-TurboID-Flag tag lentivirus plasmids were a kind gift of Steve Fuhs and Raymond Whitson (Genomics Institute of the Novartis Research Foundation, La Jolla, CA). HEK293T cells were transfected by combining 800 μL OptiMEM, 4 μg of the transgene, 2 μg psPAX2 packaging plasmid, and 1 μg of VSVG viral envelope plasmid with 21 μL of PEI 1 mg/mL. The plasmid mixture was incubated for 15 min at room temperature and added to the cells. The infection was performed as previously described in (33). Briefly, 48 h post-transfection, 9 mL of supernatant of HEK293T cells was collected, filtered through a 0.45 μm filter, combined with 1 mL of 8 mg/ml Polybrene, and added directly to target cells which were 40-50% confluent. The infection was repeated after 72 h and then transduced cells were puromycin selected (1 μg/mL) for 72 h and LHPP protein expression was evaluated by immunoblotting.

### Proximity labeling with TurboID

Stable MDA-MB-231 cells expressing LHPP-TurboID, TurboID, and the empty pLVX vector 70-90% confluent in 10 cm plates were treated with 500 μM biotin for 10 min. After treatment, protein lysates were lysed in 1X RIPA buffer, as previously described. For each condition, 300 μg of cell lysate was incubated with 25 μL of streptavidin beads (ThermoFisher Scientific, 88817) and 500 µl of 1X RIPA overnight at 4°C with agitation. The protein enrichment was performed as previously described in (34). The biotinylated proteins were eluted in 3X Laemmli buffer supplemented with 30 μL of 2 mM biotin and 20 mM DTT, boiled at 95°C for 10 min, and submitted to the Salk Institute Mass Spectrometry Core for protein identification using Tandem Mass Tag (TMT) labeling quantitative mass spectrometry.

### TMT-labeling for proteomic analysis

Samples were precipitated by methanol/chloroform and redissolved in 8 M urea/100 mM TEAB, pH 8.5. Proteins were reduced with 5 mM tris(2-carboxyethyl)phosphine hydrochloride (TCEP, Sigma-Aldrich) and alkylated with 10 mM chloroacetamide (Sigma-Aldrich). Proteins were digested overnight at 37°C in 2 M urea/100 mM TEAB, pH 8.5, with trypsin (Promega). The digested peptides were labeled with 10-plex TMT (Thermo Scientific, 90309), and pooled samples were fractionated by basic reversed phase (Thermo Scientific, 84868).

### Liquid Chromatography and Mass Spectrometry

The TMT-labeled samples were analyzed on an Orbitrap Eclipse Tribrid mass spectrometer (Thermo Scientific). Samples were injected directly onto a 25 cm, 100 μm ID column packed with BEH 1.7 μm C18 resin (Waters™). Samples were separated at a flow rate of 300 nL/min on an EasynLC 1200 (Thermo Scientific). Buffers A and B were 0.1% formic acid in water and 90% acetonitrile, respectively. A gradient of 1–10% B over 30 min, an increase to 35% B over 120 min, an increase to 100% B over 20 min, and held at 100% B for 10 min was used for a 180 min total run time. Peptides were eluted directly from the tip of the column and nanosprayed directly into the mass spectrometer by application of 2.5 kV voltage at the back of the column. The Eclipse was operated in a data dependent mode. Full MS1 scans were collected in the Orbitrap at 120k resolution. The cycle time was set to 3 s, and within this 3 s the most abundant ions per scan were selected for CID MS/MS in the ion trap. MS3 analysis with multinotch isolation (SPS3) was utilized for the detection of TMT reporter ions at 60k resolution (35). Monoisotopic precursor selection was enabled and dynamic exclusion was used with exclusion duration of 60 s.

### Proteomic analysis

Protein and peptide identification were done with Integrated Proteomics Pipeline – IP2 (Integrated Proteomics Applications). Tandem mass spectra were extracted from raw files using RawConverter (36) and searched with ProLuCID (37) against the Uniprot human database. The search space included all fully-tryptic and half-tryptic peptide candidates. Carbamidomethylation on cysteine and TMT on lysine and peptide N-term were considered as static modifications. Data were searched with 50 ppm precursor ion tolerance and 600 ppm fragment ion tolerance. Identified proteins were filtered using DTASelect (38) and utilizing a target-decoy database search strategy to control the false discovery rate to 1% at the protein level (39). Quantitative analysis of

TMT was done with Census (40) filtering reporter ions with 10 ppm mass tolerance and 0.6 isobaric purity filter.

### Immunofluorescence

Visualization of LHPP localization in MDA-MB-231 cells was performed by seeding 75,000 cells onto coverslips treated with poly-L-lysine. At 40-60% confluence, the cells were washed twice in 1X PBS and fixed in 4% PFA for 10 min. Subsequently, they were washed twice in PBS and permeabilized using 0.1% TritonX-100 in PBS and blocked in 4% BSA in PBS for 1 h. Primary antibodies were diluted in blocking buffer and incubated overnight at 4°C in a humidified chamber. The appropriate secondary antibody was diluted in blocking buffer and incubated for 1 h at room temperature. Hoechst 33342 (Invitrogen H3570, 1:5000) was used as a nuclear stain, Phalloidin Alexa Fluor 647 was used to stain F-Actin (Invitrogen A22287, 1:100) and Neutravidin 488 (Invitrogen A6374,1:350) was used to detect biotinylated proteins. The coverslips were mounted using ProLong™ Gold Antifade Mountant (Life Technologies, P6930). The primary antibodies and concentrations are as follows: mouse anti-Flag (Millipore Sigma F3165, 1:1000), rabbit anti-LHPP (Millipore Sigma, HPA009163). The secondary antibodies used were goat anti-rabbit AlexaFluor488 (Invitrogen A-11034, 1:200) and goat anti-mouse AlexaFluor546 (Invitrogen A-11030, 1:200). Confocal microscopy was performed using a Leica TCS SP8 MP microscope and Leica Application Suite X (LASX) software or a Zeiss LSM 880 Airyscan and Zen Black Software.

### ELISA

Human CCL2/MCP-1 and M-CSF/CSF1 were measured by ELISA using DuoSet kits (DY279, DY216, respectively) from R&D Systems (Minneapolis, USA) according to the manufacturer’s recommendations.

### Statistical analyses

For all measurements, a Shapiro-Wilk normality test was performed to determine if values present a Gaussian distribution, and a ROUT test to identify outliers was used. For two groups comparison, if values presented a Gaussian distribution a one-tailed unpaired Student’s t-test was used, and for values without a Gaussian distribution a Mann-Whitney test was performed. For more than two groups comparison, for normal distributed values a One-way ANOVA with Tukey’s multiple comparisons post-test was used. For more than two groups without normal distribution, a Kruskal-Wallis test with Dunn’s multiple comparison post-test was used. For scatter plots, a nonlinear curve fit was used to compare the growth rate “k” between different treatments using the extra-sum-of-squares fit test. In cell proliferation and migration curves, a Gompertz growth model was used. For tumor growth, an exponential (Malthusian) growth was used. The data are presented in the graphs with individual replicates and the standard deviation also being displayed. The normality tests, identification of outliers, group comparison tests, nonlinear curve fit, bar graphs, scatter plots, and survival curves were performed using GraphPad Prism version 8. Principal component analysis (PCA), volcano plots, heatmaps, and Gene Set Enrichment analysis were done by using the packages ggplot, pheatmap, EnhancedVolcano, and ClusterProfiler, respectively in R studio.

## Results

### LHPP levels are higher in human breast tumors than in normal tissue with the highest levels in TNBC

We initiated the study by evaluating the protein expression of LHPP in different subtypes of human breast cancer using tissue microarrays of patient breast tumors (Figure 1A). Using immunohistochemistry, we found that baseline LHPP protein expression levels are elevated compared to normal mammary tissue, with LHPP levels further increasing with the progression stage in breast tumors (Figure 1A). As LHPP is expressed mainly in the epithelial cells of the mammary glands, we observed increased LHPP levels in Ductal Carcinoma In Situ (DCIS) compared to normal tissue and tend to be at the highest level when the tumor expands and invades other tissues in Invasive Ductal Carcinoma (IDC) (Figure 1B). Furthermore, we assessed the levels of LHPP in primary breast tumor samples and normal mammary tissue from patients using immunoblotting. We found that LHPP levels were higher in breast tumors compared to normal tissue, although the difference did not reach statistical significance (Supplementary Figure S1A-B). As breast cancer is a heterogeneous disease, we investigated whether there was a particular subtype of cancer with higher expression of LHPP. Of the three major breast cancer genotypes tested we found that TNBC expressed the highest levels of LHPP compared to the ER^+^PR^+/-^HER2^+/-^ and ER^-^PR^-^HER2^+^ subtypes (Figure 1C and 1D). Notably, analysis of Kaplan-Meier Survival Curves of breast cancer patients expressing high or low levels of LHPP showed that patients expressing higher levels of LHPP have a lower probability of survival (Figure 1E). Moreover, the Kaplan-Meier survival plot of ER^+^PR^+/-^HER2^+/-^ and TNBC patients, but not ER^-^PR^-^ HER2^+^ patients, showed that patients with higher levels of LHPP have a lower probability of survival (Figure 1E-H).

**Figure 1.**
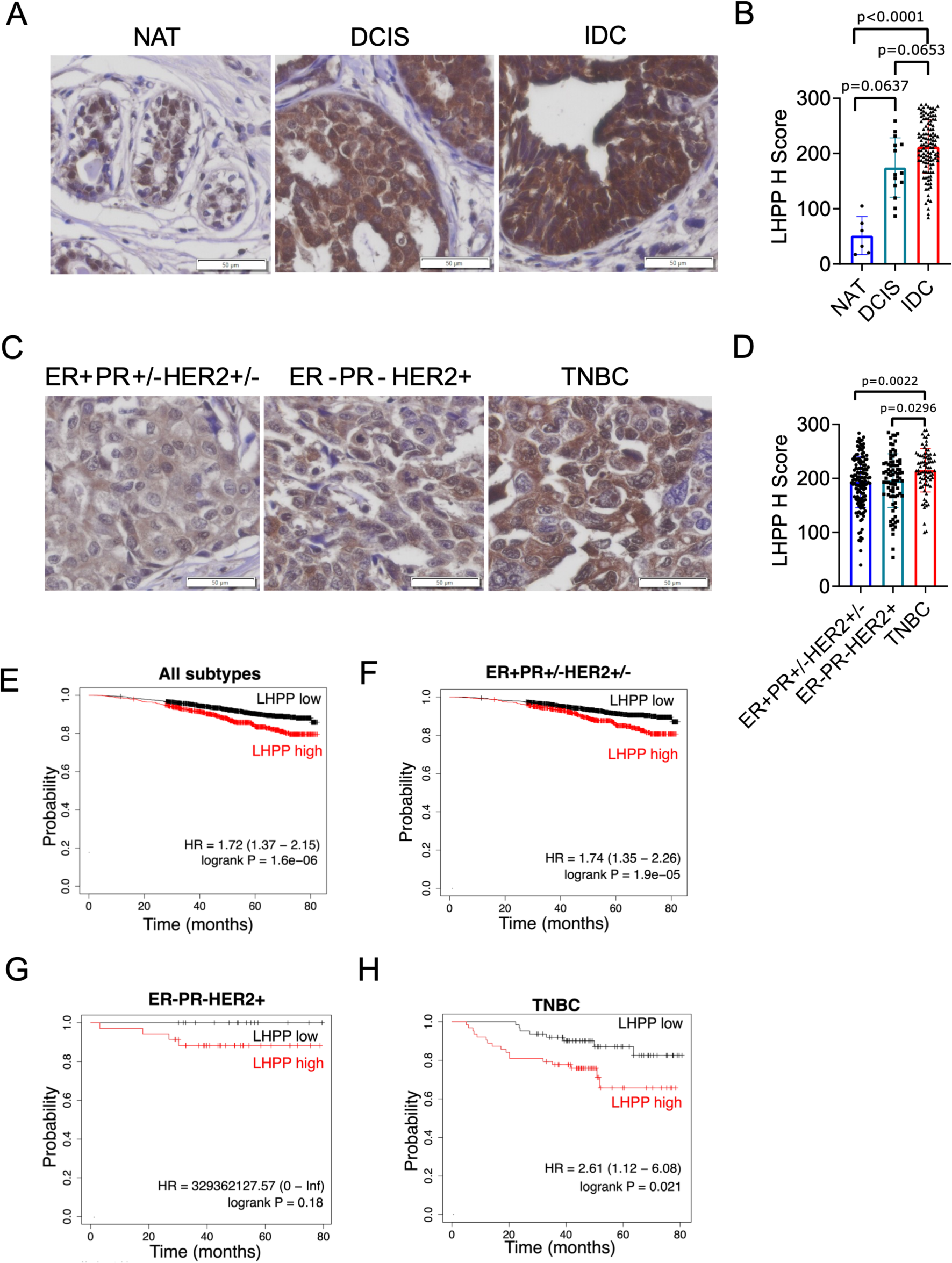
LHPP levels are higher in breast cancer compared to normal mammary tissue, with the highest levels in TNBC. **(A)** Representative images and **(B)** quantification of LHPP IHC of human breast cancer patient tissue microarray containing normal adjacent tissue (NAT) (n=3 patients in duplicate cores), ductal carcinoma in situ (DCIS) (n=7 patients in duplicate cores) and invasive ductal carcinoma (IDC) (n=60 patients in duplicate cores). Data were analyzed by the Kruskal-Wallis test with Dunn’s multiple comparison test. Data are represented as bar graphs with individual replicates ± standard deviation (SD). **(C)** Representative images and **(D)** quantification of LHPP IHC of human breast cancer patient tissue microarrays containing ER^+^PR^+/-^HER2^+/-^ subtype (n=99 patients, 40 in duplicate & in 59 single cores), ER^-^PR^-^HER2^+^ subtype (n=65 patients, 18 in duplicate & 47 in single cores), and TNBC (n=47 patients, 34 in duplicate & 13 in single cores). Data were analyzed by the Kruskal-Wallis test with Dunn’s multiple comparison test. Data are represented as bar graphs with individual replicates ± SD. **(E)** Kaplan-Meier overall survival curve of human breast cancer patients with high or low LHPP expression based on RNA levels for all breast cancer subtypes (n=2976), ER^+^PR^+/-^HER2^+/-^ subtype (n=2575 patients), ER^-^ PR^-^HER2^+^ (n=50 patients), TNBC (n=126 patients). Scale bars = 50 µm in all images.

### LHPP knockout reduces cell proliferation, cell migration, cell invasion, and spheroid formation in TNBC cells but does not affect overall phosphohistidine levels

After finding that LHPP expression was higher in breast tumors compared to normal tissue, particularly in TNBC, we used CRISPR/Cas 9 technology to knock out LHPP expression in the TNBC cell line, MDA-MB-231. We obtained two complete LHPP knockout clones: KO clone 3 (KO3) and KO clone 11 (KO11) as validated by immunoblotting (Figure 2A and 2B). By measuring the growth of the WT cells and the two KO LHPP clones, we determined that the WT cells had higher proliferation compared to both KO LHPP clones (Figure 2C). Similarly, a scratch wound assay was performed to assess the differences in cell migration between the WT and the two KO LHPP MDA-MB-231 clones. We found that WT cells had a significantly higher migration rate (Figure 2D). Next, we tested the ability of the WT and two KO clones to form spheroids in culture using ultra-low attachment plates. The two LHPP KO clones were unable to form compact spheroids as measured by their surface area (Figure 2E and 2F), which may indicate that LHPP expression affects cell-cell interactions. Finally, we also assessed if the LHPP KO clones exhibited differences in cell invasion, finding that the two LHPP KO clones invaded significantly less through transwells compared to the WT cells (Figure 2G and 2H). When we knocked out LHPP in MCF7 breast cancer cells, we observed a similar effect on cell proliferation and spheroid formation, but not on cell migration (Supplementary Figure S2A-F).

**Figure 2.**
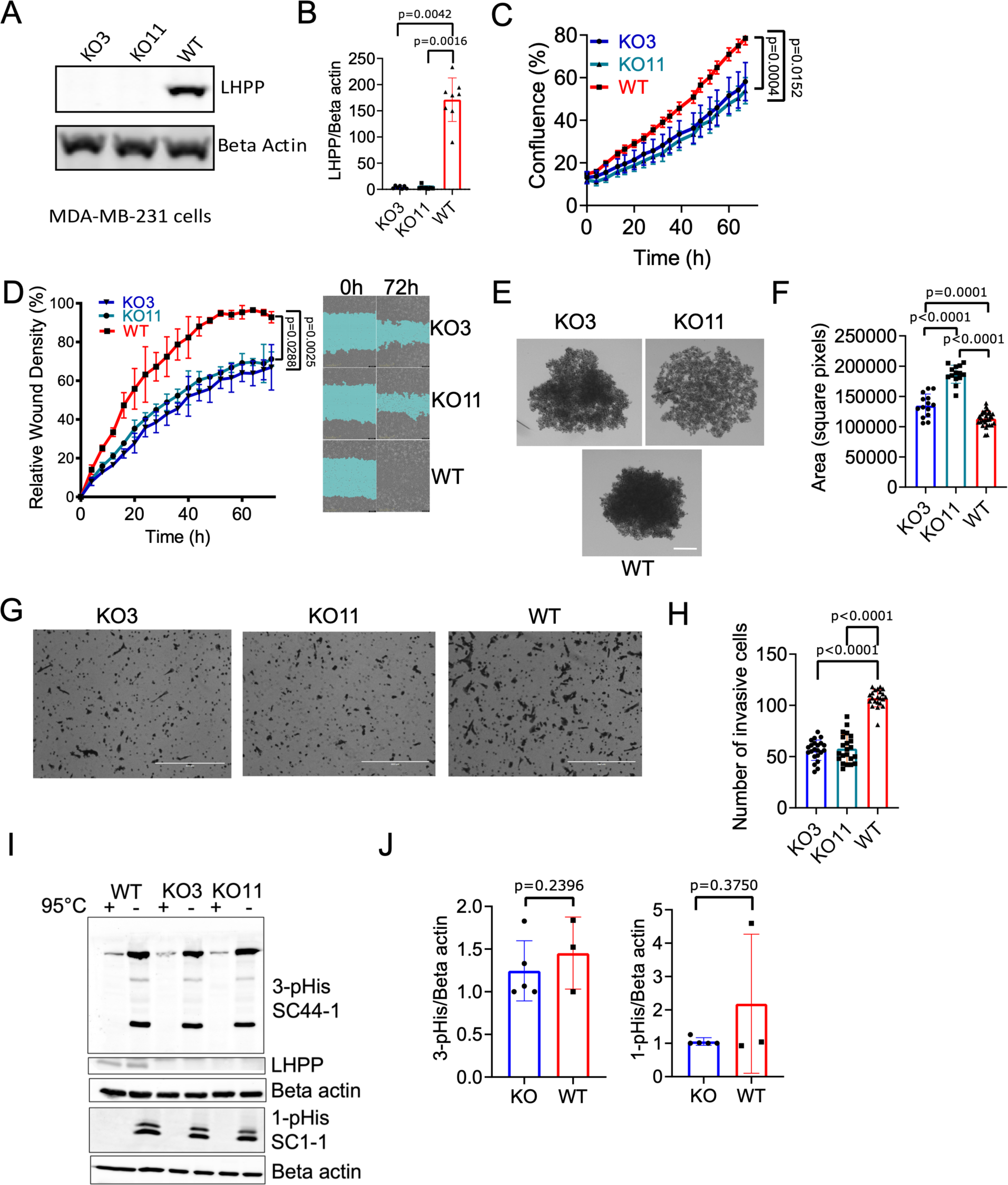
LHPP knockout decreases cell proliferation, migration, invasion, and spheroid formation but does not affect overall phosphohistidine levels in TNBC cells. **(A)** Immunoblotting and **(B)** quantification of LHPP in LHPP KO clone 3 (KO3), LHPP KO clone 11 (KO11), and WT MDA-MB-231 cells (WT) (n=4 independent experiments). Data were analyzed by the Kruskal-Wallis test with Dunn’s multiple comparison test. Data are represented as bar graphs with individual replicates ± SD. **(C)** Cell confluence percentage determined by IncuCyte software from phase-contrast images of MDA-MB-231 KO3, KO11, and WT cells (n=3 independent experiments). A non-linear fit to compare the growth rates of the curves in each condition was performed and the extra-sum-of-squares fit test was used. Gompertz growth was used as a model of cell growth. Data are represented as a scatter plot of the mean of at least 3 individual replicates ± SD. **(D)** Relative wound density measured using IncuCyte software and representative images from a scratch wound healing assay of MDA-MB-231 KO3, KO11, and WT cells (n=3 independent experiments). A nonlinear fit to compare the growth rates of the curves in each condition was performed and the extra-sum-of-squares fit test was used. Gompertz growth was used as a model of cell growth. Data are represented as a scatter plot of the mean of at least 3 individual replicates ± SD. **(E)** Representative images and **(F)** area quantification of spheroid formation assays of KO3, KO11, and WT cells, scale bar = 200 µm (n=3 independent experiments). Data were analyzed by a one-way ANOVA with Tukey’s multiple comparison test. Data are represented as bar graphs with individual replicates ± SD. **(G)** Representative images and **(H)** quantification of cell invasion assays using transwell, scale bars = 400 µm (n=3 independent experiments). Data were analyzed by the Kruskal-Wallis test with Dunn’s multiple comparison test. Data are represented as bar graphs with individual replicates ± SD. **(I)** Immunoblotting and **(J)** quantification of KO3, KO11, and WT MDA-MB-231 cells using SC44-1 3-pHis and SC1-1 1-pHis monoclonal antibodies (n=3 independent experiments). The cell lysates were boiled at 95°C as a pHis signal control, considering its instability in heat conditions. !-actin was used as a loading control. Data were analyzed by an unpaired t-test (3-pHis) and a Mann-Whitney test (1-pHis). Data are represented as bar graphs with individual replicates ± SD.

Although no direct substrate of LHPP has yet been identified, it is recognized as a phosphohistidine phosphatase, therefore we hypothesized that its protein expression level will correlate with decreased overall phosphohistidine levels in cells. To test this, we immunoblotted cell lysates from the WT and two KO clones for total 1- and 3-pHis signals using antibodies generated in house (SC1-1 and SC44-1, respectively) (3). However, there were no detectable differences between total 1- and 3-pHis expression levels in LHPP competent or deficient cells (Figure 2I). These findings suggest that either LHPP is performing a non-canonical function to regulate proliferation, migration, and invasion which is independent of regulating histidine phosphorylation, or that differences in pHis levels may require a higher resolution method of detection than immunoblotting.

### LHPP KO delays tumor growth and reduces metastasis in mice xenografts

After characterizing the phenotypic effects of LHPP expression on *in vitro* cell proliferation, migration, and invasion, we used an *in vivo* orthotopic tumor model by injecting either WT or LHPP KO3 and KO11 clones into the fourth mammary gland of NOD/SCID mice (Figure 3A). Initially, a preliminary experiment was done to find the total number of implanted cells that would yield the fastest tumor growth (Supplementary Figure S3A). It was observed that the highest tumor growth rate was achieved with 1 million MDA-MB-231 cells, compared to 0.5 million and 2 million MDA-MB-231 cells (Supplementary Figure S3A). Subsequently, 1 million MDA-MB-231 WT cells and LHPP KO cells were injected into the fourth mammary gland to examine the influence of LHPP expression on tumor formation. Tumor volumes were obtained weekly and collected at the experimental endpoint (>1000mm^3^) to assess LHPP expression levels. In addition to the primary tumor, the lungs, liver, adjacent mammary gland (second and third, #2/3 left), contralateral mammary gland (fourth, #4 right), and distant mammary gland (#2/3 right) tissues were collected to identify possible metastases. After 12 weeks, we observed potential temporal differences in tumor burden as the mice injected with WT MDA-MB-231 cells had formed tumors of ∼500 mm^3^, while the mice injected with the KO clones had a striking delay in tumor formation (Figure 3B). Mice injected with the KO clones did not form detectable tumors until 20 weeks post-injection when they began to grow at the rate of WT tumors (Supplementary Figure S3B) despite the fact that they remained LHPP-depleted at the time of collection (Figure 3J-L). This delay in tumor growth was particularly evident between KO3 and WT, with a p-value of 0.0025, and between KO11 and WT, with a p-value of 0.0019. At 20 weeks post-transplant, the percentage of mice without tumors or with tumors <1 cm^3^ was significantly higher with transplanted LHPP-deficient cells (Figure 3C).

**Figure 3.**
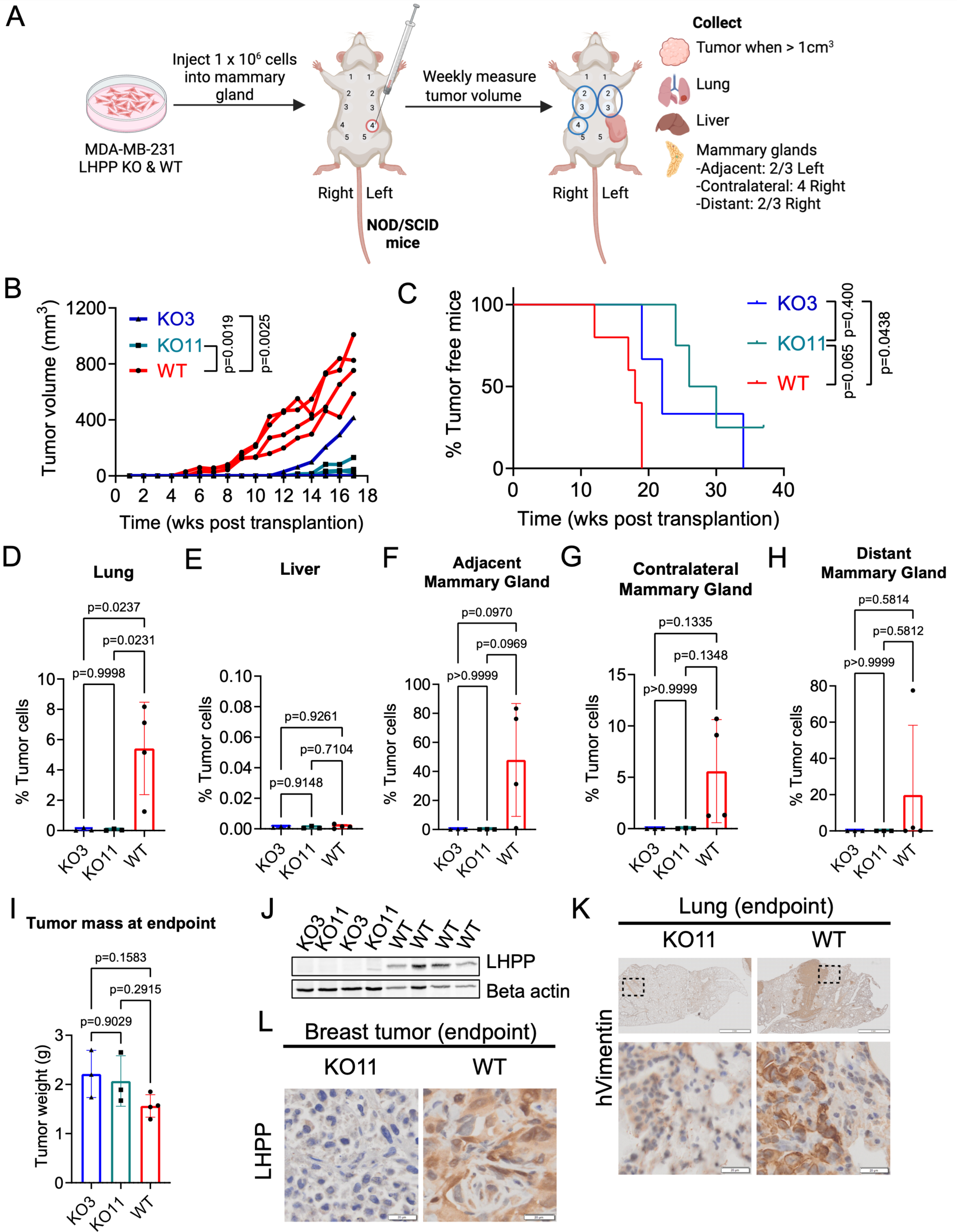
LHPP knockout delays tumor growth and decreases metastases in immunocompromised mice. **(A)** MDA-MB-231 WT and LHPP KO cells were injected into the fourth mammary gland of NOD/SCID mice; tumor growth was monitored and measured weekly until tumors reached more than 1000 mm^3^ when they were collected together with the lungs, livers, and the adjacent, contralateral, and distant mammary glands to evaluate the presence of metastases. Image created with BioRender. **(B)** Weekly tumor volume measurement until week 18 post-injection (n=4 mice). A non-linear fit to compare the growth rates of the curves in each condition was performed and the extra-sum-of-squares fit test was used. Exponential (Mathusian) growth was used as a model for tumor growth. Data are represented as a scatter plot of the tumor volume of individual replicates. **(C)** Survival curve of mice bearing tumors (n=4 mice). Mice with tumors smaller than 1000 mm^3^ volume were considered tumor-free for this curve. A Log-rank (Mantel-Cox) test was used to compare the survival curves. **(D)** Percentage of MDA-MB-231 cells detected by flow cytometry in the lung, **(E)** liver, **(F)** adjacent mammary gland, **(G)** contralateral mammary gland, **(H)** and distant mammary gland. **(I)** Tumor weight at the time of collection. Data from (D) to (I) were analyzed by a one-way ANOVA with Tukey’s multiple comparison test and n=4 mice for WT, n=3 mice for each KO clone. Data are represented as bar graphs with individual replicates ± SD. **(J)** Immunoblot of LHPP from sorted LHPP KO3, KO11, and WT tumor cells from mouse xenografts at the endpoint with !-actin used as a loading control, n=4 mice for WT, n=2 mice for each KO clone. **(K)** Representative images of human vimentin IHC from KO11 and WT MDA-MB-231 in lungs collected at endpoint from mouse xenografts bearing tumors (n=1 mouse), upper panel scale bars = 2 mm, lower panel scale bars = 20 µm. **(L)** Representative images of LHPP IHC from KO11 and WT MDA-MB-231 mouse orthotopic breast tumors (n=1 mouse), scale bars = 20 µm.

Metastasis was assessed when primary tumor reached more than 1000 mm^3^ by dissociating cells from the lung, liver, adjacent mammary glands, contralateral mammary glands, and distant mammary glands (Figure 3D-H) and analyzed by flow cytometry to check for the presence of human MDA-MB-231 cells by using specific human cell markers HLA-ABC and hCD44. The mice injected with WT MDA-MB-231 cells had metastases in all of the organs except for the liver. The mice with transplanted LHPP KO MDA-MB-231 cells had fewer metastatic events in all organs collected. Reduced metastases were not due to lower primary tumor burden since both KO and WT tumors had similar tumor weights (Figure 3I) and similar total numbers of tumor cells at the time of collection (Supplementary Figure S3C). At the experimental endpoint, we evaluated LHPP expression in MDA-MB-231 sorted cells from collected tumors, and as expected, tumors injected with WT cells maintained LHPP expression whereas the KO tumors remained LHPP-deficient (Figure 3J). We also validated LHPP expression by immunohistochemistry of the tumors collected and only the tumors with WT MDA-MB-231 cells expressed LHPP whereas the ones formed by KO LHPP clones lacked LHPP expression, confirming the results observed by immunoblotting (Figure 3L). Finally, we also validated by IHC, the presence of metastatic MDA-MB-231 WT cells in the lungs by performing a specific anti-human vimentin staining. We observed the presence of human cells in the lungs from mice injected with WT MDA-MB-231 cells but not from mice transplanted with LHPP KO MDA-MB-231 cells (Figure 3K).

### LHPP regulates the expression of genes involved in the chemokine-mediated signaling pathway

Because we did not detect striking differences in intracellular pHis levels between WT and LHPP KO cells (Figure 2I-J), and yet we found evidence of differences in tumor growth in mouse xenograft models (Figure 3B), we hypothesized that LHPP may be non-canonically regulating cell proliferation, migration, and invasion and that these effects may be reflected in LHPP-dependent transcriptional changes. To uncover the mechanisms by which LHPP is associated with these changes, we carried out RNA sequencing on WT and the LHPP KO11 clone, and performed a PCA analysis (Figure 4A) and heatmap cluster analysis (Figure 4C) to determine the similarity between the different experimental replicates of the LHPP KO11 clones and WT. Most of the differentially expressed genes (DEGs) were consistent between the experimental replicates, and 4,983 DEGs were observed between the KO11 and WT cells (Figure 4B). In the WT cells compared to the LHPP KO11 cells, 2,500 genes were found significantly downregulated, while 2,483 genes were upregulated (Supplementary Table 1). Gene set enrichment analysis of the DEGs for gene ontology biological processes showed that the LHPP depletion led to distinct profiles of genes involved in the chemokine-mediated signaling pathways, response to antigen processing, and presentation of exogenous antigen, and macrophage chemotaxis (Figure 4D). The expression of several genes involved in chemokine-mediated signaling, including *CCRL2, CXCL1, CXCL8, CXCL2, CXCR4, CCL2, CCL5, CXCR6, CMKLR1, TFF2, CCR3, ACKR3*, and *ROBO1*, was found to be increased in wild-type MDA-MB-231 cells relative to LHPP knockout cells. Similarly, we performed RNA sequencing on LHPP KO MCF7 cells and MCF7 WT cells, and found that the depletion of LHPP in MCF7 cells also significantly impacted the expression of genes related to chemotaxis (Supplementary Figure S4).

**Figure 4.**
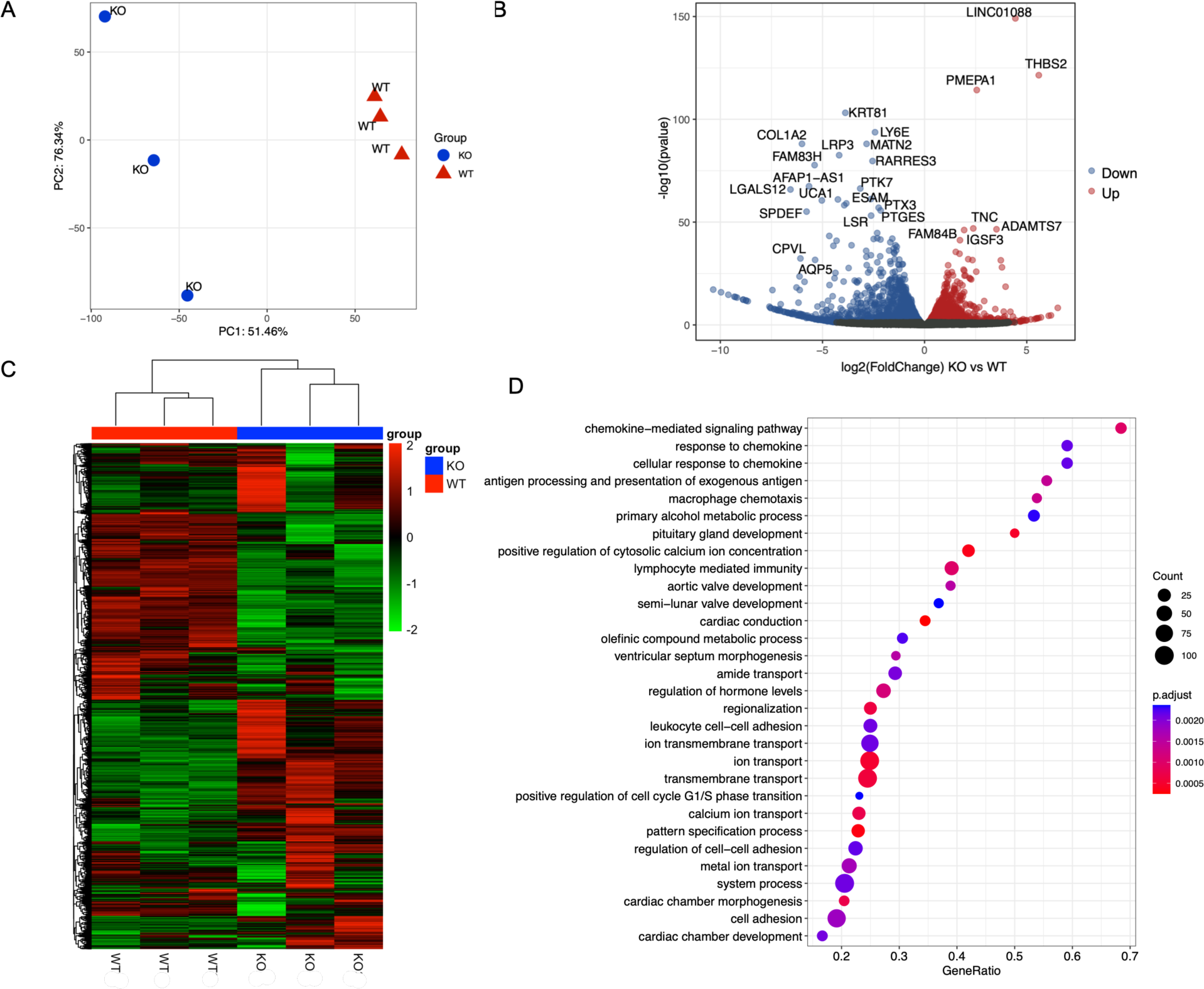
LHPP depletion in MDA-MB-231 cells affects the expression of genes involved in chemokine-mediated signaling pathways. **(A)** Principal Component Analysis of MDA-MB-231 LHPP KO11 and WT cells gene expression based on mRNA-sequencing (n=3 biological replicates). **(B)** Volcano plot of DEGs between LHPP KO11 and WT MDA-MB-231 cells (n=3 biological replicates). Blue values indicate genes that are downregulated and red values genes that are upregulated. Gray values were not statistically significant. To identify DEGs a differential gene expression analysis was done by using DESeq2 with a Benjamini-Hochberg false discovery rate and a differential gene screening threshold of log2(Fold Change) >= 1 & p-adj <= 0.05. **(C)** Heatmap of all DEGs between LHPP KO11 and WT MDA-MB-231 cells (n=3 biological replicates). FPKM values of DEGs were clustered using log2(FPKM+1) and converted to a Z-score scale. Red values indicate genes with high expression and green values indicate genes with low expression. **(D)** Dot plot from gene set enrichment analysis (Gene Ontology: biological process) of DEGs between LHPP KO11 and WT MDA-MB-231 cells (n=3 biological replicates).

### LHPP interacts with proteins involved in histone modification, mRNA processing, and actin filament organization

To further understand the mechanisms underlying LHPP’s tumorigenic effect, we performed a proximity-labeling proteomic analysis. In this assay, MDA-MB-231 cells stably expressing TurboID, a biotin ligase, fused to the C-terminus of LHPP (LHPP-TurboID) were treated with biotin, which should lead to labeling of all proteins within a 10 nm radius of an LHPP molecule, which is the radius of action of the biotin ligase (41). Biotinylated proteins were isolated by streptavidin bead affinity purification and tryptic peptides were tagged using TMT isobaric labeling and quantitative mass spectrometry (Figure 5A). Cells expressing pLVX vector only, TurboID only, or LHPP-TurboID fused protein were assessed for biotinylated proteins by performing streptavidin blotting. Both TurboID only and LHPP-TurboID fused protein can biotinylate proteins in contrast to the pLVX empty vector (Figure 5B). To validate that the LHPP-TurboID fusion protein retained phosphatase function, we performed an *in vitro* pNPP phosphatase activity assay using purified LHPP-TurboID and found that LHPP-TurboID retains phosphatase activity compared to unfused LHPP (Supplementary Figure S5). To ensure that the relatively larger size of the LHPP-TurboID fusion protein (∼65 kDa) compared to LHPP (29 kDa) did not affect its subcellular localization, we visualized the subcellular localization of LHPP-TurboID by immunofluorescence staining and determined that LHPP-TurboID localizes to both the cytoplasm and nuclei of infected cells, like the endogenous unfused LHPP (Figure 5D and 5F). We also assessed the presence of biotinylated proteins by immunofluorescence and confirmed that biotinylated proteins were only present in LHPP-TurboID expressing cells treated with biotin (Figure 5D). After LHPP-TurboID characterization by immunoblotting and immunofluorescence, we purified the biotinylated proteins from LHPP-TurboID expressing, biotin-treated MDA-MB-231 cells and identified the proteins that interacted with LHPP by TMT-labeling and mass spectrometry. We found 1457 proteins differentially labeled between LHPP-TurboID and TurboID cells (Figure 5C). After performing gene ontology biological process gene set enrichment analysis of the identified biotinylated proteins, we identified that the LHPP interactors are involved in histone modification, mRNA processing, and actin filament organization (Figure 5E). Several proteins that were enriched in LHPP-TurboID are connected with histone deacetylation, including CHD3, AKAP8L, CTBP1, CAMK2D, MTA2, SFPQ, RBM14, CHD4, HDAC1, HDAC2, GATAD2A, and GATAD2B. Additionally, proteins enriched in LHPP-TurboID related to DNA-templated transcription regulation included HDGF, MTA2, SMARCD3, BRD4, and CDK13. Regarding actin filament organization, proteins that were enriched in LHPP-TurboID were: TPM3, RAC1, CDC42, FMN1, PAK2, ROCK2, TJP1, EZR, PXN, and CTTN. In addition, we conducted LHPP proximity-labeling proteomics of in MCF7 cells and discovered that LHPP also interacts with proteins related to RNA processing in MCF7 cells (Supplementary Figure S6).

**Figure 5.**
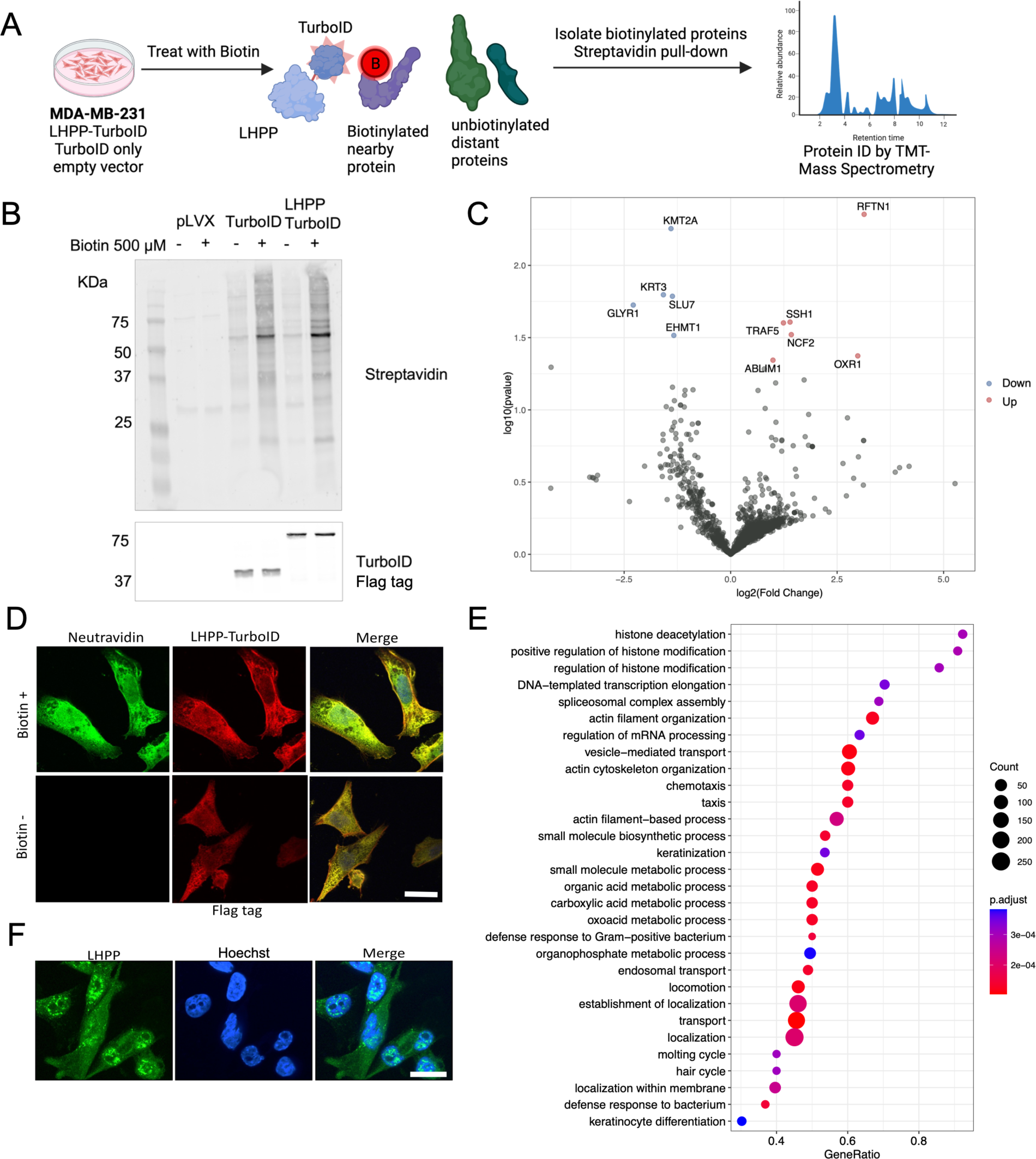
LHPP interacts with proteins involved in histone modification, mRNA processing, and actin filament organization. **(A)** MDA-MB-231 cells stably expressing LHPP-TurboID fused protein, TurboID only, or the empty vector (pLVX) were treated with biotin, and proteins in the proximity of TurboID were biotinylated. Then biotinylated proteins were isolated by affinity purification using streptavidin beads and tryptic peptides were labeled using TMT and quantified by mass spectrometry (n=3 biological replicates). Image created with BioRender. **(B)** Streptavidin immunoblot to detect biotinylated proteins in MDA-MB-231 cells treated with biotin and stably expressing the empty vector (pLVX), TurboID-Flag, and LHPP-TurboID-Flag as shown by anti-Flag tag immunoblot (n=3 biological replicates). **(C)** Volcano plot of upregulated and downregulated biotinylated proteins between MDA-MB-231 cells expressing TurboID and LHPP-TurboID (n=3 biological replicates). Blue values indicate genes that are downregulated and red values genes that are upregulated. Gray values were not statistically significant. A Student’s t-test was performed to determine the statistical significance of labeled proteins between samples. **(D)** Representative image of immunofluorescence of biotinylated proteins stained with neutravidin of MDA-MB-231 expressing LHPP-TurboID-Flag fused protein and treated with biotin as shown with immunofluorescence of Flag tag, scale bar = 20 µm (n=10 cells, 2 independent experiments). **(E)** Dot plot from gene set enrichment analysis (Gene Ontology: biological process) of biotinylated proteins between MDA-MB-231 cells expressing TurboID and LHPP-TurboID (n=3 biological replicates). **(F)** Representative image of immunofluorescence of endogenous LHPP in MDA-MB-231 cells. Hoechst 33342 was used to stain the nuclei, scale bar = 20 µm (n=10 cells, 3 independent experiments).

We found that actin filament organization, actin cytoskeleton organization, and actin-filament-based processes were enriched in the proteomic proximity-labeling analysis (Figure 5E) and regulation of cell-cell adhesion and cell adhesion were processes prominent in the DEGs between KO11 LHPP and WT cells (Figure 4D). Moreover, we found that the cytoskeleton-associated proteins such as fascin actin-bundling protein 1 (FSCN1), tropomyosin 3 (TPM3), receptor for activated C kinase 1 (RACK1), and p21-activated kinase 2 (PAK2) genes were upregulated in WT cells compared to LHPP KO11 cells (Supplementary Figure S7A). Most of the proteins encoded by these genes were also identified and enriched in LHPP-TurboID-expressing cells (Supplementary Table 2). Therefore, we stained for F-actin using phalloidin and found that LHPP WT cells have more filopodia than LHPP KO11 cells (Supplementary Figure S7B and S7C). Also, KO11 LHPP cells appeared to have more intracellular stress fibers, although this difference was not quantified.

### LHPP regulates the recruitment of macrophages to the tumor microenvironment

After gaining insight into possible LHPP oncogenic mechanisms in TNBC cells, we validated some of these targets using tumor mouse xenografts. The RNAseq DEGs data was enriched for chemokine-mediated signaling pathway, response to chemokine, cellular response to chemokine, and macrophage chemotaxis biological processes (Figure 4D). The proteomic proximity-labeling analysis also showed enrichment for chemotaxis and taxis (Figure 5E). Moreover, secretion of the macrophage-recruiting chemokines CCL2 and CSF1 were significantly attenuated in the LHPP KO11 clone versus WT MDA-MB-231 cells (Figure 6A). The protein levels of CCL2 and CSF1 were confirmed to be diminished in the supernatant of LHPP KO11 MDA-MB-231 cells relative to the supernatant of WT MDA-MB-231 cells by ELISA (Figure 6B). These findings led us to investigate whether macrophage recruitment within the mouse xenograft tumors was affected by LHPP expression in tumor cells. Specific staining to detect F4/80^+^ macrophages revealed that the KO11 cell tumors had fewer intratumoral macrophages as compared to the WT tumors. These differences were especially noticeable at early time points when KO tumors were significantly smaller than the WT ones (Figure 6C, left panel). In the KO11 cell tumors, most of the macrophages were in the peritumoral area; however, in the WT cell tumors, there was a stronger presence of macrophages inside the tumor. In contrast, specific staining for cell proliferation by Ki67 showed that both KO11 LHPP and WT cells achieved a proliferative state from week 1 after transplantation (Figure 6C, right panel). This indicates that tumor-infiltrating macrophages rather than intrinsic tumor cell proliferation are likely promoting tumor growth in the WT mouse xenografts after transplantation. Together, these results validate some of our transcriptomics and proximity-labeling observations and suggest that the differences in tumor growth between the WT and LHPP KO cell mouse xenografts could be due to LHPP-dependent recruitment of macrophages to the tumor microenvironment.

**Figure 6.**
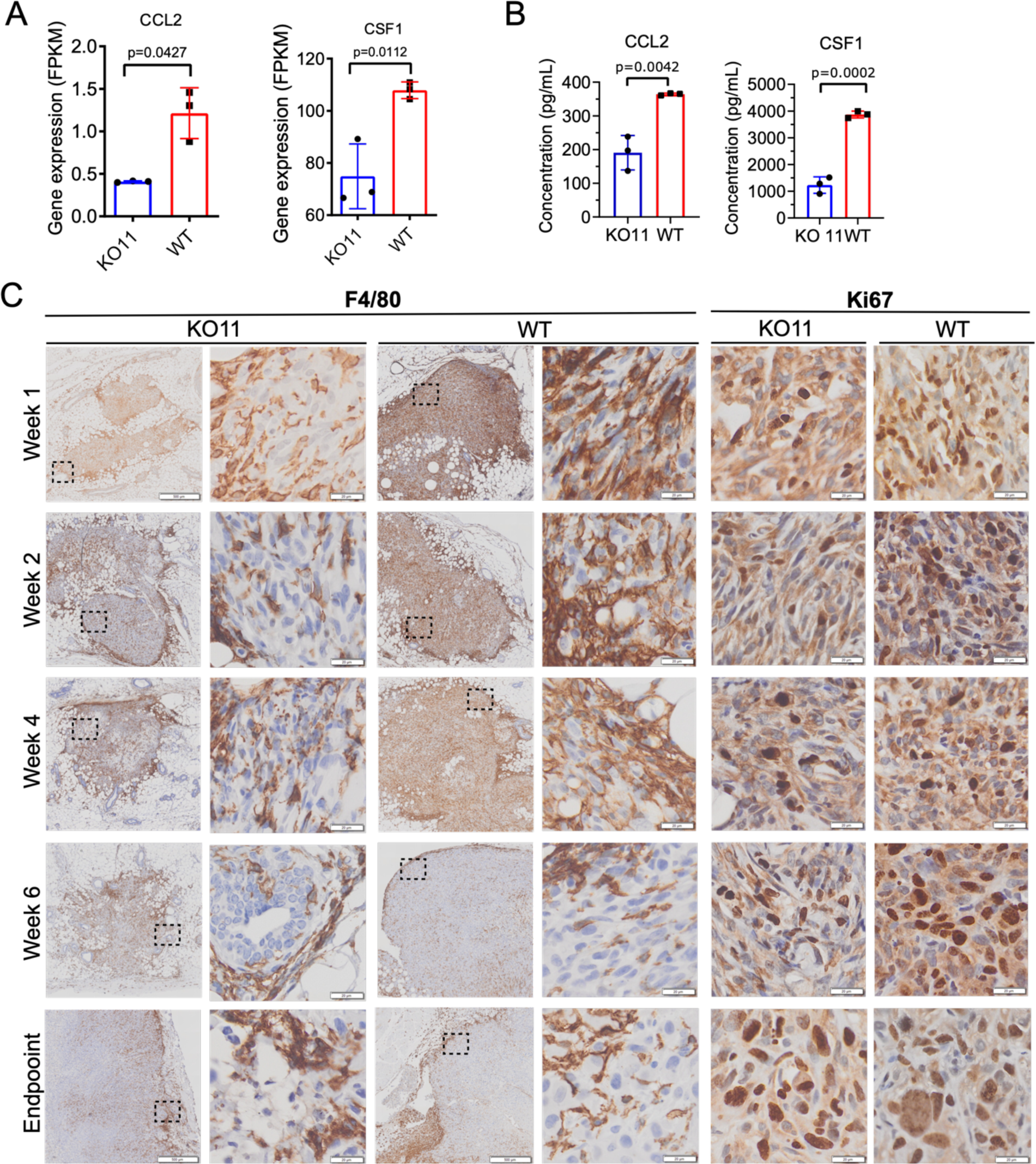
LHPP expression in tumor cells recruits macrophages to the tumor microenvironment. **(A)** Gene expression of macrophage recruiting factors CCL2 and CSF1 in MDA-MB-231 LHPP KO11 and MDA-MB-231 WT cells as measured by mRNA-sequencing (n=3 biological replicates). Data were analyzed by an unpaired t-test. Data are represented as bar graphs with individual replicates ± SD. **(B)** Quantification of CCL2 and CSF1 protein in LHPP KO11 and WT MDA-MB-231 cells supernatants measured by ELISA (n=3 biological replicates). Data were analyzed by an unpaired t-test. Data are represented as bar graphs with individual replicates ± SD. **(C)** Representative images of IHC of F4/80 and Ki67 to detect macrophages and proliferating cells, respectively, in mouse tumor xenografts from LHPP KO11 and WT MDA-MB-231 cells (n=1 mouse per time point). Lower magnification scale bars = 500 µm, higher magnification scale bars = 20 µm.

## Discussion

In this study, we investigated the role of the LHPP phosphohistidine phosphatase in triple-negative breast cancer (TNBC) and its impact on tumor growth and metastasis. We found that LHPP expression is significantly increased in TNBC compared to normal breast tissue, and, based on LHPP knockout studies, we conclude that loss of LHPP expression delays TNBC cell tumor growth and reduces formation of lung metastasis *in vivo*. These findings suggest that LHPP may play an important role in the aggressiveness and metastatic potential of TNBC. By exploring the molecular mechanisms through which LHPP regulates tumor growth and metastasis in TNBC using mRNA sequencing and proximity-labeling proteomics, we found that LHPP expression in TNBC cells led to increased recruitment of macrophages into the tumor microenvironment.

The results of several previous studies of the role of LHPP in different cancers, including hepatocellular carcinoma, renal cancer and gastric cancer are consistent with LHPP having a role as a tumor suppressor. Our study, however, suggests a distinct role for LHPP in breast cancer, specifically in TNBC, where we have shown that LHPP is required for tumor growth and metastasis by examining human patient samples, human TNBC cells, and a mouse xenograft TNBC model. These results indicate that LHPP might have a dual role as a tumor suppressor and tumor promoter, depending on the tissue type, highlighting the importance of understanding LHPP in a context-specific manner. A protein having a dual role as an inhibitor or promoter of tumor progression is not unprecedented and has also been reported for the NME1 histidine kinase, which in pediatric neuroblastoma patients, in contrast to adult tumors, is associated with poor prognosis and a more aggressive phenotype (5). Several other tumor suppressors have been reported to have dual roles as oncogenes depending on their cellular context (42).

Although the majority of studies have proposed a tumor-suppressive role for LHPP, such as in hepatocellullar carcinoma (4), RNAseq databases, such as GEPIA2 (43) suggest that there might be an oncogenic role in 14 cancer types where LHPP RNA expression is higher in tumor tissues than in normal tissues - breast cancer, lymphoid neoplasm, diffuse large B-cell lymphoma, kidney renal papillary cell carcinoma, acute myeloid leukemia, lower-grade glioma, liver hepatocellular carcinoma, lung adenocarcinoma, lung squamous cell carcinoma, ovarian serous cystadenocarcinoma, pancreatic adenocarcinoma, stomach adenocarcinoma, thyroid carcinoma, thymoma, and uterine corpus endometrial carcinoma. However, LHPP protein levels in these cancer types have not been confirmed in cancer cell lines, mouse models, or patient samples, and further work would be required to determine whether LHPP might have a tumor-promoting role in these cancers. In addition, LHPP levels were higher in the tumor cells compared to normal tissues in a mouse PDAC model (44), suggesting a tumor-promoting role for LHPP in this cancer type, in contrast to prior reports for PDAC (20, 45).

Two other phosphohistidine phosphatases, PHPT1 (12–15) and PGAM5 (16, 17), have also been implicated in promoting tumorigenesis. However, unlike PHPT1 and PGAM5, structural analysis suggests that the LHPP catalytic site is not accessible to large proteins, and may only be accessible to small phosphate-containing molecules, like other phosphatases in the HAD family (46), most of which only act on small molecules and not phosphoproteins (46). Loss of LHPP expression in MDA-MB-231 (Figure 2I-J) and MCF7 cells (Supplementary Figure S2G-H) did not lead to detectable changes in phosphohistidine protein levels by immunoblotting, although the anti-pHis antibodies used in these will not detect all possible pHis proteins in these cells (47), and further studies are required. Importantly, it is necessary to conduct experiments in which a catalytically dead mutant of LHPP is expressed in the LHPP KO cells to determine the significance of its catalytic function for the newly discovered tumor-promoting function.

Our transcriptomics and proximity-labeling of proteins results are consistent with LHPP being important for small-molecule metabolism. The transcriptomics data showed that depletion of LHPP affects the expression of genes involved in the primary alcohol metabolic processes and the olefinic compound metabolic process (Figure 4D). Additionally, our proximity-labeling proteomics data showed that LHPP is in proximity to proteins involved in small-molecule metabolism, organic acid metabolism, and organophosphate metabolism (Figure 5E). These findings provide a list of potential metabolic processes to investigate for the involvement of LHPP, and potential phosphate-containing small molecules to test as LHPP substrates. The precise manner in which LHPP regulates gene expression, and whether its function is nuclear or cytoplasmic also remain to be elucidated.

Published reports indicate that LHPP expression mostly affects the PI3K/AKT signaling pathway, and in some cancers, such as liver cancer, it also affects TGF-β/phospho-Smad signaling (27). Our proximity-labeling proteomics revealed some proteins involved in TGF-β signaling, such as PAK2 and ROCK2 (Supplementary Table 2). However, no extensive studies of LHPP’s interacting partners have been reported, which would be essential to advance our understanding of its function. One study has identified NME1 and NME2 as substrates for LHPP in neurons (48), and another showed that LHPP induces ubiquitin-mediated degradation of PKM2 in glioblastoma, although PKM2 was not an LHPP phosphotarget (26). Neither of these proteins was regulated by LHPP expression as detected by immunoblotting or by *in vitro* biolayer interferometry with purified proteins (data not shown), even though proximity labeling identified NME1 and NME2 as interactors (Supplementary Table 2). Our list of LHPP’s potential interactors will be a basis for future studies to identify LHPP substrates and expand the knowledge of its molecular mechanisms and involvement in other biological processes.

The significant role that LHPP plays in the development of metastases may be attributed to its ability to modulate the organization of the cytoskeleton and regulate the expression of proteases that are involved in cell invasion. Our proximity-labeling and next-generation sequencing data are consistent with LHPP being important for actin cytoskeleton organization. The observed differences in filopodia formation and cytoskeleton organization (Figure S7B and S7C) might explain the less invasive phenotype of LHPP KO MDA-MB-231 cells both *in vitro* and *in vivo*. Interestingly, other phosphatases related to LHPP, such as PHPT1 and Eya3, have also been shown to modulate cytoskeletal organization (12, 49, 50). LHPP has also been shown to play a significant role in regulating expression of specific matrix metalloproteinases MMP-2 and MMP-9 in cervical cancer cells (19) and MMP-7 and MMP-9 in liver cancer cells (51), proteases that are responsible for breaking down the extracellular matrix, and enabling tumor cells to invade other tissues. Furthermore, our transcriptomics analysis of LHPP KO MDA-MB-231 cells revealed reduced expression levels of the ADAM proteases, including ADAMTS9, ADAMTS15, and ADAMTS1 compared to LHPP WT MDA-MB-231 cells, suggesting another possible explanation of LHPP’s crucial role in metastases.

Our differential gene expression analysis also hinted at an involvement of LHPP in macrophage chemotaxis through regulation of expression of macrophage-recruiting factors CCL2 and CSF1 in TNBC cells. CCL2 and CSF1 were downregulated in LHPP KO MDA-MB-231 cells, and in the first weeks after transplantation, orthotopic LHPP KO mouse tumors presented fewer F4/80+ tumor-infiltrating macrophages, which are important for tumor initiation and progression. High levels of tumor-associated macrophages in TNBC tumors are correlated with poor prognosis, tumor progression, and a heightened risk of metastasis (52–54). LHPP’s association with immune cell responses has previously only been documented in genome-wide association studies (GWAS) (55). This may be a novel function of LHPP and it would be important to explore how LHPP affects the chemokine-mediated signaling of macrophages.

In summary, we have uncovered a tumor-promoting role of LHPP in breast cancer, which is distinct from its proposed tumor suppressor role in other cancer types, and adds to our understanding of how this pHis phosphatase can play distinct roles depending on the cancer context.

## Acknowledgments

The authors would like to thank Liliana Zamora and Ana Peterson for technical assistance in histological analysis and tumor measurements. Additionally, we thank Stephen Fuhs and Raymond Whitson for providing the LHPP-TurboID plasmids. We thank all Hunter lab and Wahl lab members for their support and discussions. JR is a Pew Latin American Fellow, supported by The Pew Charitable Trusts, and also by a Curebound Discovery Grant (#626773). QVM was supported by a Swiss National Foundation Mobility Fellowship (P2ELP3_199760) and a Catharina Foundation Fellowship. JN was supported by Salk Woman in Science Award from the Salk Institute. ANM was supported by an NIH T32 Cancer Training Grant from the Salk Institute. KTY was supported in part by American Cancer Society IRG Grant # IRG-19-230-48-IRG and UC San Diego Moores Cancer Center, Specialized Cancer Center Support Grant NIH/NCI P30CA023100. Studies conducted in the Wahl lab were funded by grants from the Breast Cancer Research Foundation, The Freeberg Foundation, and the National Institutes of Health (*NIH/NCI (R35 CA197687) and NCI Core Grant CA014195).* T.H. is a Frank and Else Schilling American Cancer Society Professor and holds the Renato Dulbecco Chair in Cancer Research. T.H. is supported by an NIH National Cancer Institute R35 (5 R35 CA242443) award. Research funding for this project was also provided by a Curebound Discovery Grant (#626773) and a Salk Cancer Center Pilot Grant. This work was supported by the Mass Spectrometry Core of the Salk Institute with funding from NIH-NCI CCSG: P30 CA01495, NIH-NlA San Diego Nathan Shock Center P30 AG068635, an NIH S10 award for metabolic instrumentation: S10 OD021815, and the Helmsley Center for Genomic Medicine. We thank J. Diedrich and A. M. Pinto for mass spectrometry technical support. This research was made possible with the support of the Flow Cytometry Core Facility of the Salk Institute (RRID: SCR_014839) with funding from NIH-NCI CCSG: P30 CA01495 and Shared Instrumentation Grant S10-OD023689. This project received support from the Waitt Advanced Biophotonics Core Facility of the Salk Institute with funding from NIH-NCI Cancer Center Support Grant (CCSG): P30 CA01495, NIH-NlA San Diego Nathan Shock Center P30 AG068635, and the Waitt Foundation.

