## Supplementary Method and Figures for "LHPP expression in triple-negative breast cancer promotes tumor growth and metastasis by modulating the tumor microenvironment"

### Supplementary methods

*LHPP immunoblot of human patient simples:*
50 mg of normal and breast cancer fresh frozen samples in liquid nitrogen were provided by the Biorepository and Tissue Technology Shared Resources from Moores Cancer Center at the University of California San Diego (UCSD). All de-identified human tissue samples were previously in existence. Breast tumor samples were from patients diagnosed with triple-negative breast cancer, stage II-III with post-neoadjuvant treatment. The normal tissue was obtained from patients undergoing reduction mammaplasty. The Multi-sample BioSpec BioPulverizer (Biospec, BS:59012MS) was utilized to pulverize the frozen tissue. The powdered tissue was weighed (20-40 mg) and 10 μL of RIPA 1X pH 8.8 were added per 1 mg of tissue. The VWR® 200 Homogenizer (VWR, 10032-336) was employed to homogenize the tissue, which was then incubated at 4°C for at least 1 h and 30 min. The lysates were subsequently cleared and quantified using the same conditions as for whole cell lysates, as previously described. Immunoblot analysis was conducted using the same protocol as for whole cell lysates. The rabbit anti-LHPP antibody (Sigma Millipore ZRB1050, 1:1000) was used to detect LHPP, and Revert™ 700 Total Protein Stain (LI-COR, 926-11021) was used to normalize LHPP expression in human patient tissue samples.

*Cell proliferation, cell migration, and spheroid formation of MCF7 cells:*
MCF7 breast cancer cells were obtained from the Salk Cancer Center collection and cultured under the same conditions as MDA-MB-231 cells. MCF7 cells that had undergone CRISPR/Cas9-mediated knockout of LHPP using the same guide RNA as for the MDA-MB-231 cell LHPP KOs, as well as their corresponding non-edited control cells, were utilized in a series of assays, including cell proliferation, cell migration, and spheroid formation. The assays were conducted like those employed for the MDA-MB-231 cell line. The MCF7 LHPP KO cells generated were from a single clone.

*mRNA sequencing mRNA-seq) in MCF7 cells*:
The analysis of MCF7 and LHPP KO MCF7 cells was conducted using the same methodology as the analysis of MDA-MB-231 cells for mRNA-seq.

*Generation of LHPP WT and LHPP catalytic mutant cell lines:*
The LHPP-Flag tag wild type (WT) and LHPP D17N, D214N catalytic double mutant (CM) sequences were cloned by designing gBlocks™gene fragments (Integrated DNA Technologies) with the respective LHPP sequence and inserting it into the pLVX vector using the In-Fusion® Snap Assembly Master Mix (Takara). The transduction of the plasmids and selection were done in the same manner as described for LHPP-TurboID stable cell lines.

*p-nitrophenyl phosphate (pNPP) assay:*
In the presence of LHPP pNPP is converted into p-nitrophenolate ion which absorbs at 405 nm. A 100 μl reaction volume was prepared with TBS pH7.4, with a final concentration of 25 mM pNPP, 5 mM $MgCl_2$, and 500 μM of purified LHPP-TurboID-Flag tag, LHPP-Flag tag, and LHPP catalytic mutant (CM)-Flag tag. LHPP CM was made by the following mutations on LHPP amino acid sequence: D17N and D214N. LHPP-TurboID and unfused LHPP were purified from MDA-MB-231 cells stably expressing LHPP-TurboID-Flag tag, LHPP-Flag tag, and LHPP CM-Flag tag using anti-Flag beads (Millipore Sigma, A2220) and eluted with 3X-Flag peptide solution (Millipore Sigma, F4799). The reaction was incubated at 37°C, and the absorbance at 405 nm was read at 0, 14, 81, 142, and 204 min.

*Proximity labeling with TurboID of LHPP in MCF7 cells*:

MCF7 cells stably expressing LHPP-TurboID-Flag, TurboID-Flag, and the empty pLVX vector were used. These cells were analyzed in the same manner as for MDA-MB-231 cells for proximity labeling.

*F-actin staining and filopodia quantification analysis:*

LHPP KO11 and WT MDA-MB-231 cells were grown on coverslips and fixed as mentioned previously. Phalloidin Alexa Fluor 647 was used to stain F-actin (Invitrogen, A22287, 1:100), and Hoechst 33342 (Invitrogen, H3570, 1:5000) was used to stain F-actin and the nuclei respectively. The cells were imaged on a Leica TCS SP8 MP microscope and the number of filopodia, and their length were quantified by using the draw line function on ImageJ.

### Supplementary Figures

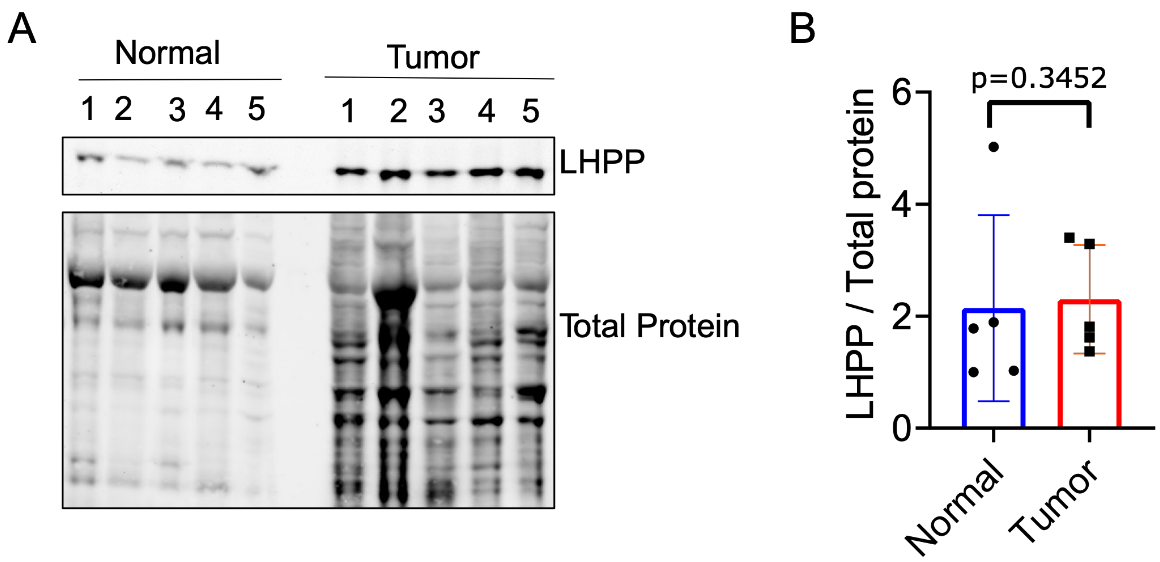


**Figure S1. Levels of LHPP are elevated in human primary breast tumors as compared to normal breast tissue. (A)** Immunoblot and **(B)** quantification of LHPP and total protein levels in human primary breast tumors (n=5 patients) and normal breast (n=5 patients) tissue samples. Data were analyzed by a Mann-Whitney test and are represented as bar graphs with individual replicates ± standard deviation (SD)

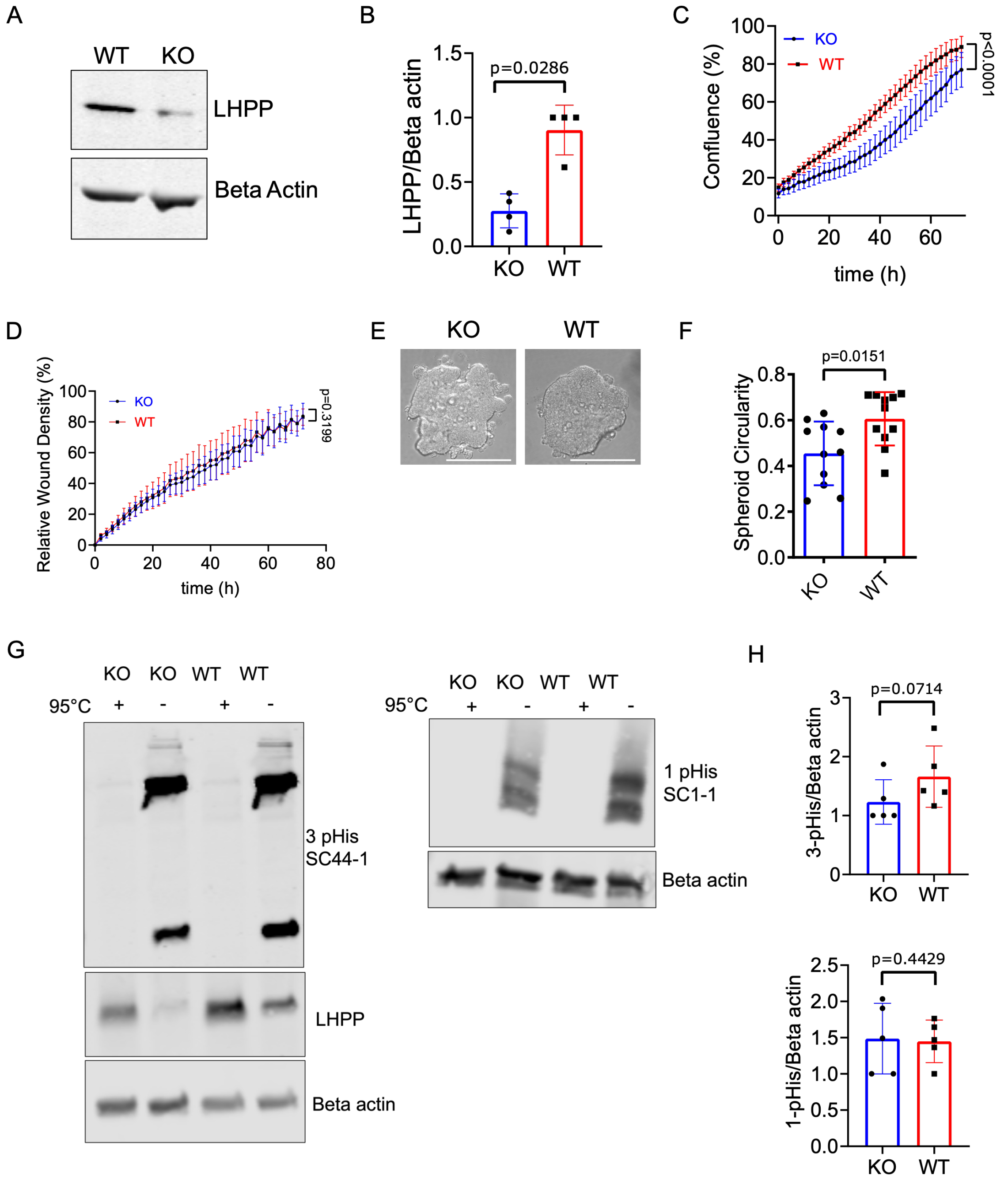


**Figure S2**. **LHPP knockout decreases cell proliferation, and spheroid formation in MCF7 cells but does not affect cell migration and phosphohistidine protein levels**. **(A)** Immunoblotting and **(B)** quantification of LHPP in LHPP KO, and WT MCF7 cells (n=4 independent experiments). Data were analyzed by a Mann-Whitney test and are represented as bar graphs with individual replicates ± SD. **(C)** Cell confluence percentage determined by IncuCyte software from phase-contrast images of LHPP KO, and WT MCF7 cells (n=3 independent experiments). A non-linear fit to compare the growth rates of the curves in each condition was performed and the extra-sum-of-squares fit test was used.

Data are represented as a scatter plot of the mean of at least 3 individual replicates ± SD. **(D)** Relative wound density was measured using IncuCyte software from a scratch wound healing assay of LHPP KO, and MCF7 WT cells (n=3 independent experiments). A non-linear fit to compare the growth rates of the curves in each condition was performed and the extra-sum-of-squares fit test was used. Data are represented as a scatter plot of the mean of at least 3 individual replicates ± SD. **(E)** Representative images and **(F)** spheroid circularity quantification of spheroid formation assays with LHPP KO and WT MCF7 cells, scale bar = 400 µm (n=3 independent experiments). Data were analyzed by a Mann-Whitney test and are represented as bar graphs with individual replicates ± SD. **(G)** Immunoblotting and **(J)** quantification of LHPP KO and WT MCF7 cells using SC44-1 3-pHis and SC1-1 1-pHis monoclonal antibodies to monitor 3-pHis and 1-pHis protein levels, respectively (n=3 independent experiments). The cell lysates were boiled at 95°C as a pHis signal control, considering its instability in heat conditions. Endogenous LHPP was detected by blotting with an LHPP antibody. Beta-actin was used as a loading control. Data were analyzed by a Mann-Whitney test (3-pHis) and an unpaired t-test (1-pHis). Data are represented as bar graphs with individual replicates ± SD.

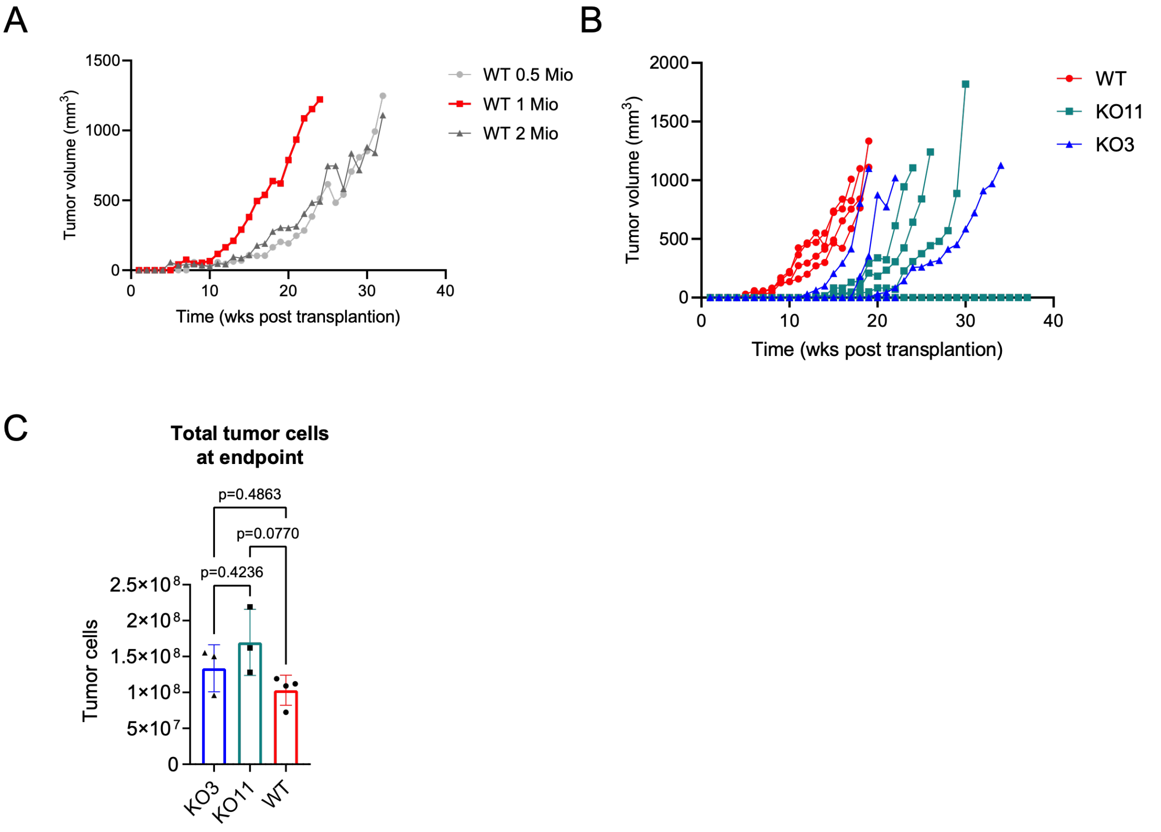


**Figure S3. LHPP knockout delays MDA-MB-231 tumor growth and decreases metastases in immunocompromised mice. (A)** Weekly tumor volume measurement of 0.5, 1, or 2 x $10^6$ MDA-MB-231 WT cells injected into the fourth mammary gland of NOD/SCID mice (n=1 mouse). Data are represented as a scatter plot of the tumor volume of individual replicates. **(B)** Weekly tumor volume measurement of the NOD/SCID mice injected with 1 x$10^6$ WT, LHPP KO11, and LHPP KO3 MDA-MB-231 cells until they reached the endpoint volume (n=4 mice). Data are represented as a scatter plot of the tumor volume of individual replicates. **(C)** Total number of tumor cells at the time of collection. Data were analyzed by a one-way ANOVA with Tukey's multiple comparison test and n=4 mice for WT, n=3 mice for each KO clone. Data are represented as bar graphs with individual replicates ± SD.

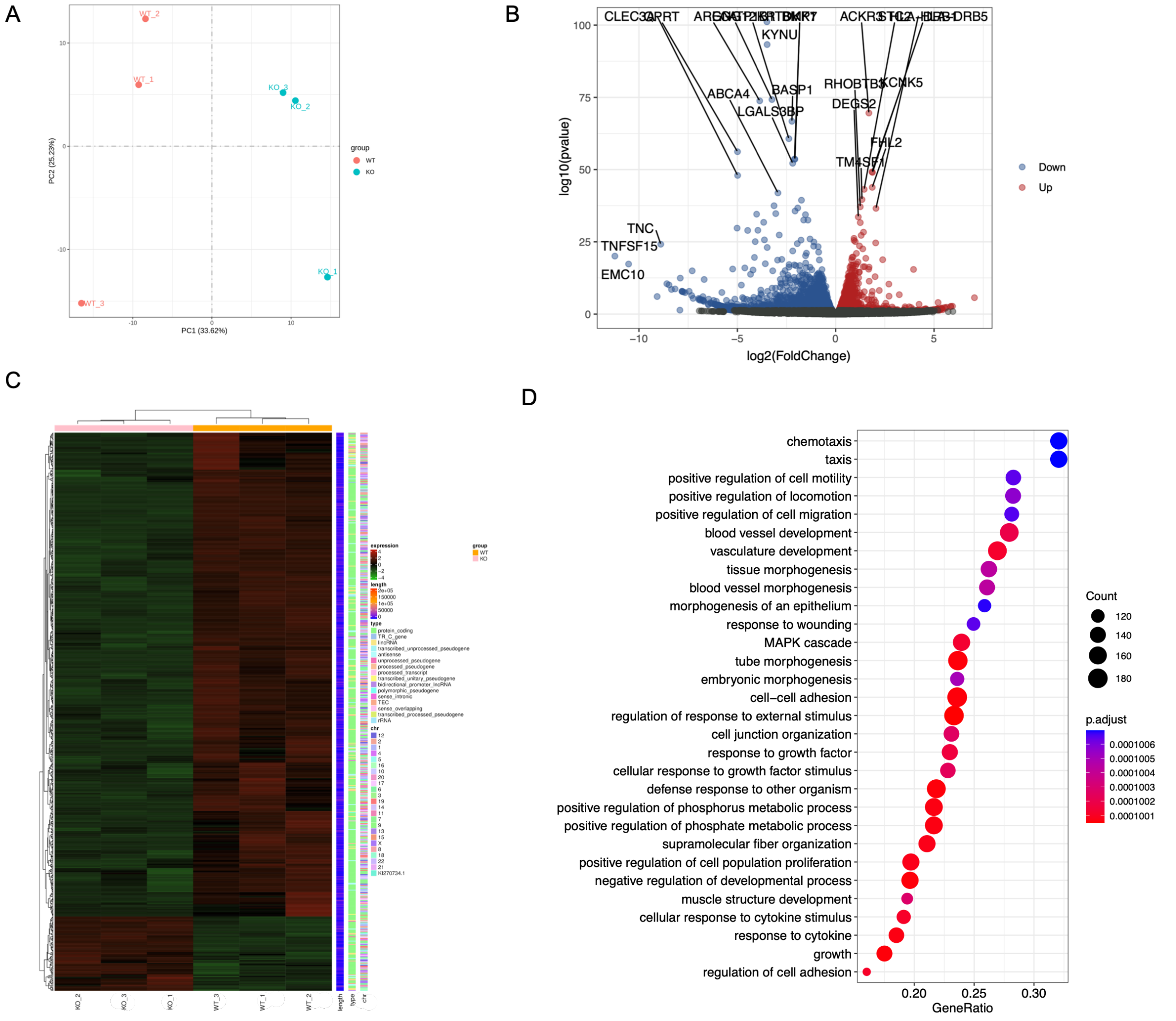


**Figure S4. LHPP depletion in MCF7 cells affects the expression of genes involved in chemotaxis and cell migration**. **(A)** Principal Component Analysis of LHPP KO and WT MCF7 cell gene expression based on mRNA-sequencing (n=3 biological replicates). **(B)** Volcano plot of differentially expressed genes (DEGs) between LHPP KO and WT MCF7 cells (n=3 biological replicates). Blue values indicate genes that are downregulated and red values genes that are upregulated. Gray values were not statistically significant. To identify DEGs a differential gene expression analysis was done by using DESeq2 with a Benjamini-Hochberg false discovery rate and a differential gene screening threshold of log2(Fold Change) >= 1 & p-adj <= 0.05. **(C)** Heatmap of all DEGs between LHPP KO and WT MCF7 cells (n=3 biological replicates). FPKM values of DEGs were clustered using log2(FPKM+1) and converted to a Z-score scale. Red values indicate genes with high expression and green values indicate genes with low expression. **(D)** Dot plot from gene set enrichment analysis

(Gene Ontology: biological process) of DEGs between LHPP KO and WT MCF7 cells (n=3 biological replicates).

A

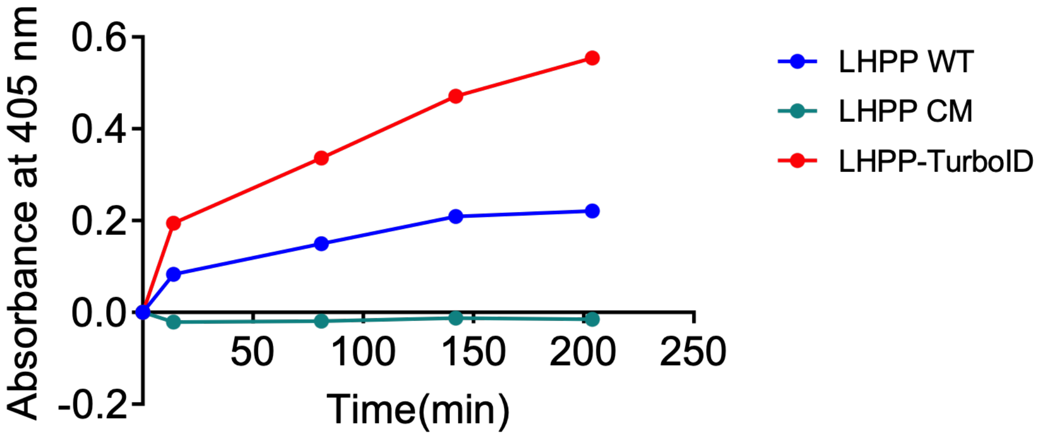


**Figure S5. LHPP-TurboID fused protein retains its phosphatase activity measured by pNPP assay. (A)** Scatter plot of the phosphatase activity of LHPP-TurboID fused protein, LHPP catalytic mutant (D17A, D214N), and WT purified proteins from MDA-MB-231 cells as measured by pNPP assay.

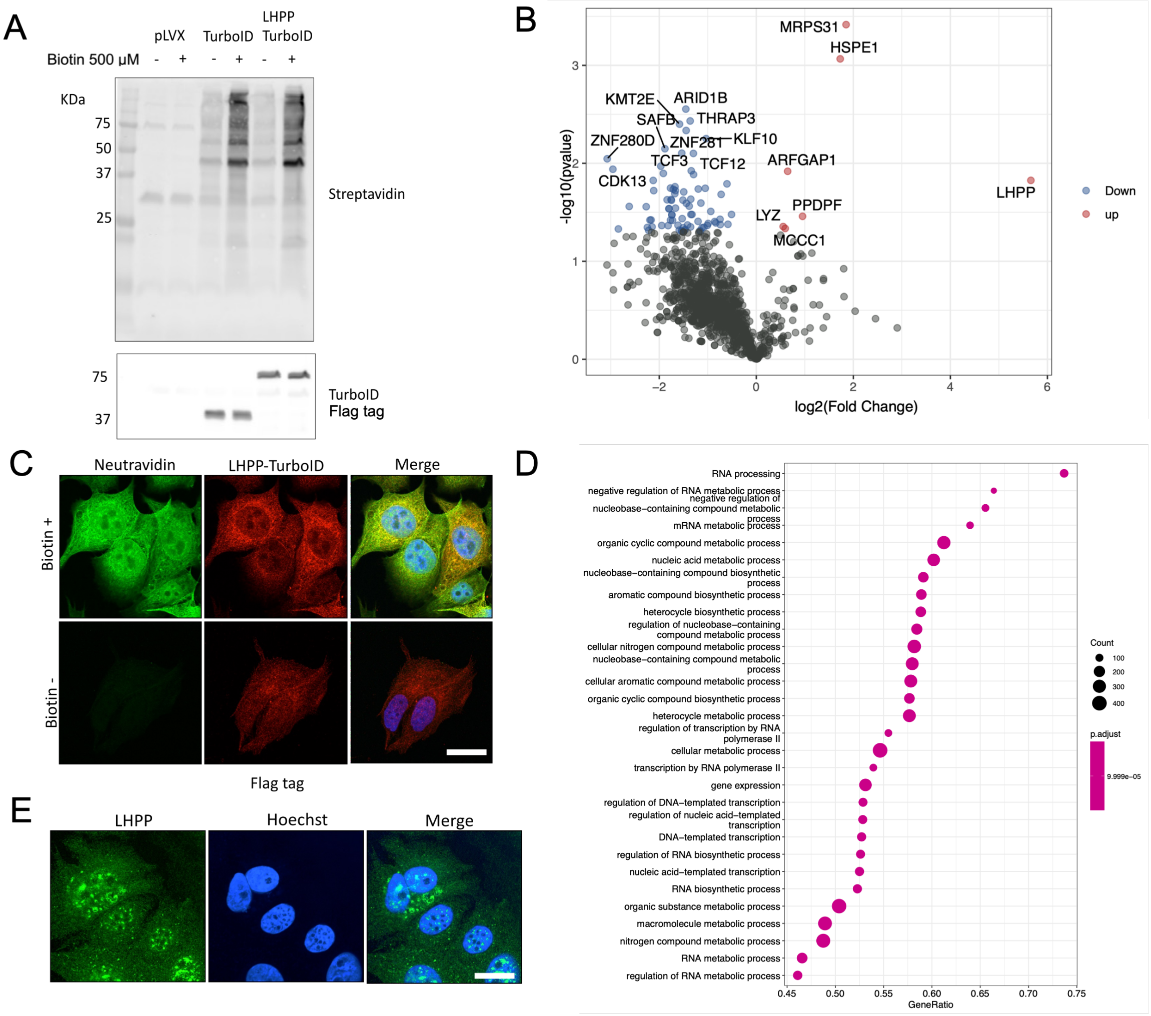


**Figure S6. LHPP interacts with proteins involved in mRNA processing and nucleotide metabolism in MCF7 cells.** MCF7 cells stably expressing Flag-tagged LHPP-TurboID fused protein, Flag-tagged TurboID only, or the empty vector (pLVX) were treated with biotin, and proteins in proximity of TurboID were biotinylated. Then biotinylated proteins were isolated by pull-down with streptavidin beads and identified by TMT-Mass Spectrometry. **(A)** Streptavidin blot to detect biotinylated proteins in MCF7 cells treated with biotin and stably expressing the empty vector (pLVX), TurboID-Flag, and LHPP-TurboID-Flag as shown by anti-Flag tag immunoblot (n=3 biological replicates). **(B)** Volcano plot of upregulated and downregulated biotinylated proteins between MCF7 cells expressing TurboID and LHPP-TurboID (n=3 biological replicates). Blue values indicate genes that are downregulated and red values genes that are upregulated. Gray values were not statistically significant. A Student's t-test was performed to determine the statistical significance of labeled proteins between samples. **(C)** Representative image of immunofluorescence of biotinylated proteins stained with neutravidin of MCF7 expressing LHPP-TurboID-Flag fused protein and treated with biotin as shown with immunofluorescence of Flag tag, scale bar = 20 μm (n= 10 cells). **(D)** Dot plot from gene set enrichment analysis (Gene Ontology: biological process) of biotinylated proteins between MCF7 cells expressing TurboID and LHPP-TurboID (n=3 biological replicates). **(E)** Representative image of immunofluorescence of endogenous

LHPP in MCF7 cells. Hoechst 33342 was used to stain the nuclei, scale bar = 20 μm (n=10 cells, 3 independent experiments).

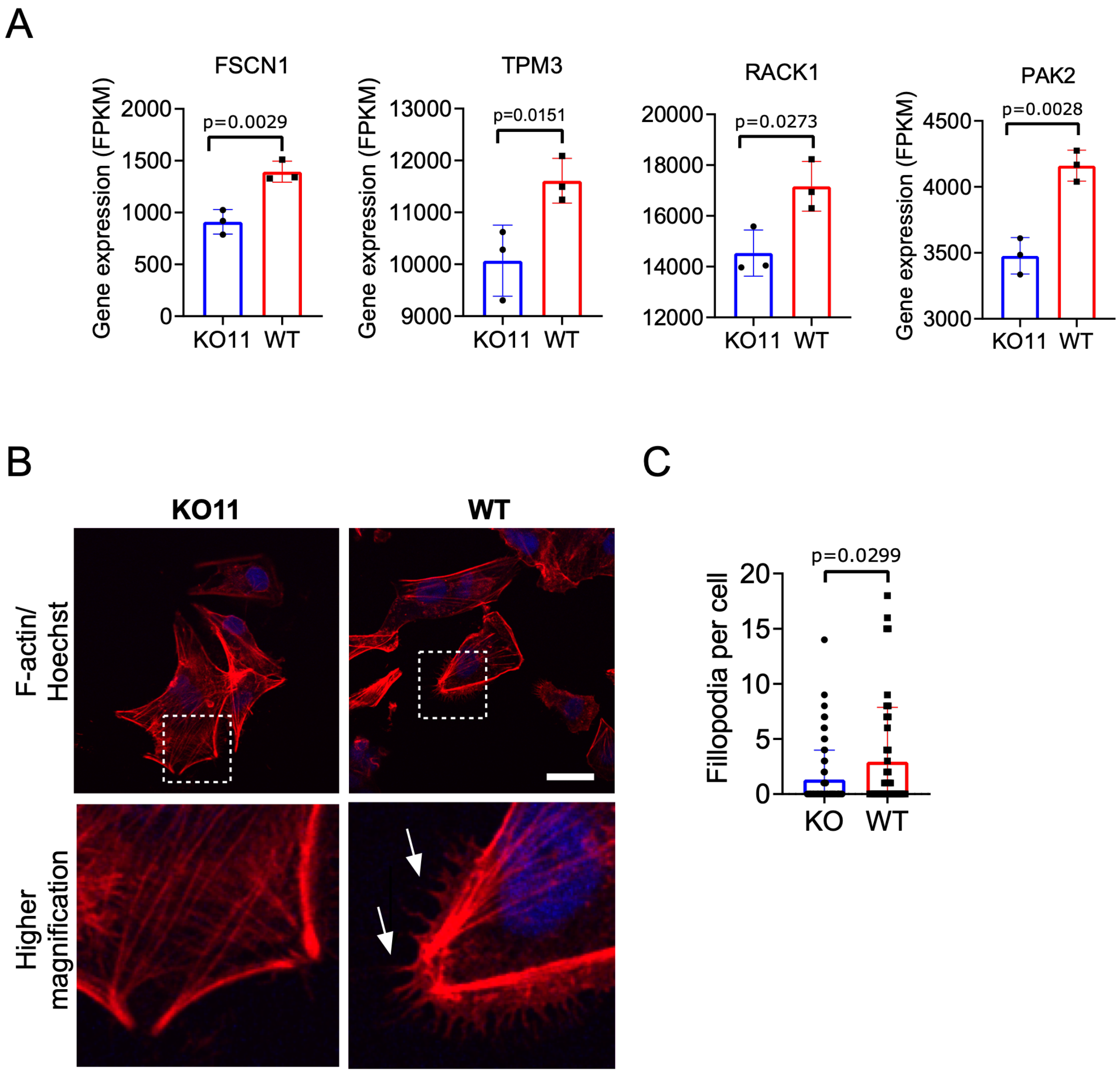


**Figure S7. LHPP knockout decreases filopodia formation in MDA-MB-231 cells. (A)** Gene expression of FSCN1, TPM3 RACK1, and PAK2 in LHPP KO11 and WT MDA-MB-231 cells as measured by mRNA-sequencing (n=3 biological replicates). Data were analyzed by an unpaired t-test. Data are represented as bar graphs with individual replicates ± SD. **(B)** F-actin staining by phalloidin of LHPP KO11 and WT MDA-MB-231 cells. Hoechst 33342 was used to stain nuclei, arrows point to filopodia structures, scale bar = 20 μm. **(C)** Filopodia per cell quantification of phalloidin stained LHPP KO11 (n=80 cells) and WT MDA-MB-231 cells (n=44 cells).
